# Activation of MrgprA3 and MrgprC11 on bladder-innervating afferents induces peripheral and central hypersensitivity to bladder distension

**DOI:** 10.1101/2020.12.18.423548

**Authors:** Luke Grundy, Ashlee Caldwell, Sonia Garcia-Caraballo, David Grundy, Nick J. Spencer, Xinzhong Dong, Joel Castro, Andrea M. Harrington, Stuart M. Brierley

## Abstract

Understanding the sensory mechanisms innervating the bladder is paramount to developing efficacious treatments for chronic bladder hypersensitivity conditions. The contribution of Mas-gene-related G protein-coupled receptors (Mrgpr) to bladder signalling is currently unknown. Here we show in mice with single-cell RT-PCR that sub-populations of dorsal root ganglion (DRG) neurons innervating the mouse bladder express *MrgprA3* (14%) and *MrgprC11 (38%)*, either individually or in combination, with high levels of co-expression with *Trpv1* (81-89%). Calcium imaging studies demonstrated MrgprA3 and MrgprC11 agonists (chloroquine, BAM8-22 and neuropeptide FF) activated sub-populations of bladder-innervating DRG neurons, showing functional evidence of co-expression between MrgprA3, MrgprC11 and TRPV1. In *ex vivo* bladder-nerve preparations chloroquine, BAM8-22 and neuropeptide FF all evoked mechanical hypersensitivity in sub-populations (20-41%) of bladder afferents. These effects were absent in recordings from *Mrgpr-clusterΔ*^*−/−*^ mice. *In vitro* whole-cell patch clamp recordings showed that application of an MrgprA3/C11 agonist cocktail induced neuronal hyper-excitability in 44% of bladder-innervating DRG neurons. Finally, *in vivo* instillation of an MrgprA3/C11 agonist cocktail into the bladder of wild-type mice induced a significant activation of dorsal horn neurons within the lumbosacral spinal cord, as quantified by pERK-immunoreactivity. This MrgprA3/C11 agonist-induced activation was particularly apparent within the superficial dorsal horn and the sacral parasympathetic nuclei of wild-type, but not *Mrgpr-clusterΔ*^*−/−*^ mice. This study demonstrates, for the first time, functional expression of MrgprA3 and MrgprC11 in bladder afferents. Activation of these receptors is not required for normal bladder function but does trigger hypersensitivity to distension, a critically valuable factor for therapeutic target development.

**Significance statement:** Determining how bladder afferents become sensitized is the first step in finding effective treatments for common urological disorders such as overactive bladder and interstitial cystitis/bladder pain syndrome. Here we show that two of the key receptors, MrgprA3 and MrgprC11, that mediate itch from the skin are also expressed on afferents innervating the bladder. Activation of these receptors results in sensitization of bladder afferents, resulting in sensory signals being sent into the spinal cord that prematurely indicate bladder fullness. Targeting bladder afferents expressing MrgprA3 or MrgprC11 and preventing their sensitisation may provide a novel approach for treating overactive bladder and interstitial cystitis/bladder pain syndrome.

## INTRODUCTION

Patients suffering from common urological disorders such as overactive bladder (OAB) and interstitial cystitis/bladder pain syndrome (IC/BPS) experience chronic urinary urgency, frequency, and discomfort or pain (Abrams et al., 2002; Homma et al., 2020). Hypersensitivity of the peripheral afferents innervating the bladder are considered a key component in the underlying symptomology of these conditions (de Groat and Yoshimura, 2009; de Groat et al., 2015; Grundy et al., 2018a). Numerous nociceptive ion channels and receptors, including TRPV1 and P2X3, have been identified as contributing to bladder afferent hypersensitivity, and have been subsequently explored in order to develop novel treatments for bladder disorders (Daly et al., 2007; de Groat and Yoshimura, 2009; DeBerry et al., 2014; Yoshimura et al., 2014; Grundy et al., 2018d; Grundy et al., 2018b; Andersson, 2019). However, other mechanisms are also likely involved. In this regard, cues can be taken from sensations that originate from the skin, whereby itch and pain are commonly experienced protective sensations that occur in response to irritant or noxious stimuli (Steinhoff et al., 2018). Both of these processes require activation and sensitisation of sensory afferents that project from the skin into the dorsal horn of the spinal cord where they provide input to the central nervous system (Liu and Ji, 2013; LaMotte et al., 2014; Schmelz, 2015; Lee et al., 2016). Interestingly, itch can be mediated by histamine release from immune cells, which activate histamine receptors on sensory afferents (Shim and Oh, 2008), whilst several histamine-independent itch mechanisms have also been described. These include those dependent on Mas related G-protein coupled receptors (Mrgprs) (Tominaga and Takamori, 2014; Zhu et al., 2017), a large subfamily of GPCRs predominantly expressed in the dorsal root ganglia (DRG) (Dong et al., 2001; Lembo et al., 2002). In particular, MrgprA3 has been identified as the receptor responsible for itch associated with the antimalarial drug chloroquine (Liu et al., 2009), whilst MrgprC11 activation via the endogenous peptide agonist bovine adrenal medulla 8-22 (BAM8-22) or mast cell mediator neuropeptide FF (NPFF), a dual MrgprC11 and A4 agonist (Liu et al., 2009) also initiate pruritus and scratching (Lee et al., 2008; Sikand et al., 2011).

Traditionally these mechanisms have been considered unique to the skin, however, pruritogenic-like mechanisms are being increasingly identified in the viscera. The same histamine receptors that mediate pruritus have been explored in a number of visceral organs, including the gastrointestinal tract and bladder, where histamine has been found to activate and induce hypersensitivity of visceral afferents (Kreis et al., 1998; Wouters et al., 2016; Grundy et al., 2020). More recently, MrgprC11 and MrgprA3 have been localised in subpopulations of colon-innervating afferents (Castro et al., 2019; Van Remoortel et al., 2019), as well as vagal afferents innervating the lungs (Han et al., 2018). Activation of MrgprA3 and MrgprC11 expressed on colonic afferents results in colonic hypersensitivity, enhanced abdominal pain and altered animal behaviours (Castro et al., 2019; Van Remoortel et al., 2019), effects that were significantly increased in an animal model of chronic visceral hypersensitivity (Castro et al., 2019). Given the relative similarities in the extrinsic sensory afferent innervation of the colon and bladder, including the ability to encode a variety of nociceptive and non-nociceptive stimuli (Brierley et al., 2004; Zagorodnyuk et al., 2006; Xu and Gebhart, 2008; Grundy et al., 2019a), we sought to explore the expression of key Mrgprs in the bladder and their role in mediating neuronal excitability and bladder sensory signalling.

Here, we show that MrgprC11, MrgprA3, and to a much smaller extent MrgprD are expressed in sub-populations of bladder-innervating DRG neurons. Agonists of MrgprC11 and MrgprA3 cause pronounced afferent hypersensitivity to bladder distension *ex vivo*, activate and evoke hyperexcitability of bladder-innervating DRG neurons *in vitro*, as well as enhancing activation of neurons within the dorsal horn of the spinal cord *in vivo*.

## METHODS

### Ethics and animals

In this study, 12-to 18-week-old male and female C57BL/6J mice from an in-house breeding colony at the South Australian Health and Medical Research Institute (SAHMRI) were used. Mice were acquired from an in-house C57BL/6J breeding programme (JAX strain #000664; originally purchased from The Jackson Laboratory; breeding barn MP14; Bar Harbor, ME) and then bred within the specific and opportunistic pathogen-free animal care facility at SAHMRI. A genetically modified mouse strain lacking a cluster of MRGPR genes (*Mrgpr-clusterΔ*^*−/−*^), including MrgprA3 and MrgprC11, was utilised for some experiments (Liu et al., 2009). The *Mrgpr-clusterΔ*^*−/−*^ mice were a generous gift from Prof Xinzhong Dong (Howard Hughes Medical Institute, Johns Hopkins University, Baltimore, Maryland, USA) then bred in-house at SAHMRI as per the C57BL/6J mice. All experiments were approved by and performed in accordance with the approval of the SAHMRI Animal Ethics Committee (SAM195 and SAM190). Mice were group housed (up to 5 mice per cage) within individual ventilated cages filled with chip coarse dust-free aspen bedding (Cat#–ASPJMAEBCA; PuraBed, Niederglatt, Switzerland), with free access to LabDiet JL Rat and Mouse/Auto6F chow (Cat #5K52, St. Louis, MO) and autoclaved reverse osmosis water. Cages were stored within a temperature-controlled environment of 22°C and a 12-hour light/12-hour dark cycle on individual ventilated cage racks. Experiment sample sizes are detailed in the corresponding figure legends.

### *Ex vivo* bladder afferent nerve recordings

Nerve recording was performed using a previously described *ex vivo* model (Grundy et al., 2018d; Grundy et al., 2018b; Grundy et al., 2018c; Grundy et al., 2019b; Konthapakdee et al., 2019; Grundy et al., 2020). Mice were humanely killed via CO_2_ inhalation, and the entire lower abdomen was removed and submerged in a modified organ bath under continual perfusion with gassed (95% O_2_ and 5% CO_2_) Krebs-bicarbonate solution (composition in mmol/L: 118.4 NaCl, 24.9 NaHCO_3_, 1.9 CaCl_2_, 1.2 MgSO_4_, 4.7 KCl, 1.2 KH_2_PO_4_, 11.7 glucose) at 35°C. The bladder, urethra, and ureters were exposed by removing excess tissue. Ureters were tied with 4-0 perma-hand silk (Ethicon, #LA53G). The bladder was catheterised (PE 50 tubing) through the urethra and connected to a syringe pump (NE-1000) to allow a controlled fill rate of 30 μL/min with saline (NaCl, 0.9%). A second catheter was inserted through the dome of the bladder, secured with silk, and connected to a pressure transducer (NL108T2; Digitimer) to enable intravesical pressure recording during graded distension. Pelvic nerves, isolated from all other nerve fibres between the pelvic ganglia and the spinal cord, were dissected into fine multiunit branches and a single branch was placed within a sealed glass pipette containing a microelectrode (WPI) attached to a Neurolog headstage (NL100AK; Digitimer). Nerve activity was amplified (NL104), filtered (NL 125/126, bandpass 50–5,000 Hz, Neurolog; Digitimer), and digitised (CED 1401; Cambridge Electronic Design, Cambridge, UK) to a PC for offline analysis using Spike2 software (Cambridge Electronic Design, Cambridge, UK). The number of action potentials crossing a pre-set threshold at twice the background electrical noise, was determined per second to quantify the afferent response. Single-unit analysis was performed offline by matching individual spike waveforms through linear interpolation using Spike2 version 5.18 software. A single unit was deemed to be responsive to an agonist if a greater than 20% increase in neuronal excitability (impulses/second across all distension pressures 0-30mmHg (AUC)) was detected.

#### Afferent recording experimental protocols

At the start of each afferent recording experiment, control bladder distensions were performed with intravesical infusion of saline (NaCl, 0.9%) at a rate of 30 μL/min to a maximum pressure of 30 mmHg at 10 min intervals to assess the viability of the preparation and reproducibility of the intravesical pressure and neuronal responses to distension. The volume in the bladder was extrapolated from the known fill rate (30 μL/min) and the time taken (s) to reach maximum pressure (30 mmHg). Compliance was determined by plotting intravesical pressure against the calculated volume. After a stable baseline was maintained, the saline in the infusion ump was replaced by either BAM8-22 (20μM; Tocris Bioscience; #1763), Chloroquine (100μM; Sigma, #PHR1258-1G), Neuropeptide FF (NPFF: 50μM; Tocris Bioscience; #3137), or an ‘itch cocktail’ containing all three agonists combined (BAM8-22: 20μM, Chloroquine: 100μM and NPFF: 50μM). All experimental compounds were dissolved in dH_2_O to form stock solutions which were frozen at −80°C and defrosted immediately before being used for each experiment. The single nerve bundle isolated and inserted into the glass electrode during the dissection process was contained within the recording electrode for the entire experiment, allowing multiunit afferent nerve recordings to be performed and comparisons between afferent firing rates to be compared in the same nerve fibres during intra-bladder incubation with saline and Mrgpr agonists.

### Retrograde tracing from the bladder

A small, aseptic, abdominal incision was made in anaesthetised (2% – 4% isoflurane in oxygen) mice. A 5μL Hamilton syringe attached to a 30-gauge needle was used to inject Cholera Toxin subunit B conjugated to AlexaFluor 488 (CTB-488, 0.5 % diluted in 0.1 M phosphate buffer saline [PBS] pH 7.4; ThermoFisher Scientific) at three sites into the bladder wall (3 μL/injection) (Grundy et al., 2018a; Grundy et al., 2018c; Grundy et al., 2020). To prevent injection of CTB into the bladder lumen, the needle was inserted subserosally, parallel with the bladder muscle. The abdominal incision was then sutured closed and analgesic (Buprenorphine (Temvet); 0.1 mg/kg; Troy Laboratories Pty Ltd, APVMA #67612) and antibiotic (Amoxicillin; 50 mg/kg; Amoxil, AUSTR11137) administered subcutaneously as mice regained consciousness. Following laparotomy, mice were individually housed and allowed to recover. After 4 days, mice were humanely euthanised and the lumbosacral (LS; L5-S1) dorsal root ganglia (DRG) removed for subsequent isolation and culture of the neurons to visualise CTB-labelled bladder innervating neurons amongst the DRG neurons.

### mRNA expression analysis in DRG, isolated urothelial cells and bladder mucosal and detrusor layers

#### Tissue collection and urothelial cell isolation

16-to 18-week-old mice were humanely euthanised via CO_2_ inhalation and the bladder and lumbosacral (L5-S1) dorsal root ganglia (DRG) removed. DRG were frozen in dry ice in pairs by spinal level (L5, L6, S1) and stored at −80°C for RNA extraction and quantitative reverse-transcription polymerase chain reaction (RT-qPCR). The bladder was opened, stretched out and pinned flat, urothelial side up. For tissue RT-qPCR, the mucosal and detrusor layers were gently peeled apart, frozen in dry ice and stored at −80°C for RNA extraction and RT-qPCR. For urothelial cell RT-qPCR, urothelial cells were isolated as performed previously (Grundy et al., 2018d; Grundy et al., 2018b; Grundy et al., 2020). Following a rinse in sterile PBS, the bladder was incubated for 3 hours at room temperature in DMEM containing 2.5 mg/mL dispase (GIBCO, ThermoFisher Scientific, #17105041), 10% foetal calf serum, and 1 μM/mL HEPES (Sigma, #H3375, pH 7.0). A blunt scalpel was used to collect cells by gentle scraping of the urothelium. Cells were dissociated in 0.025% trypsin EDTA (GIBCO, ThermoFisher Scientific, #25200072) at 37°C in 5% CO_2_ for 10 min with gentle intermittent trituration. The cell suspension was then added to DMEM containing 10% foetal calf serum to deactivate the trypsin before centrifugation (15 min, 1500 rpm, 4°C). Following aspiration of supernatant, cells were resuspended in keratinocyte serum-free media (KSFM; Invitrogen, #17005042) and cell count and viability were determined using the Countess Automated Cell Counter (Invitrogen, ThermoFisher Scientific). Cells were pelleted, frozen in dry ice and stored at −80°C for RNA extraction and RT-qPCR.

#### RNA extraction

RNA was extracted using the PureLink RNA Micro kit (Invitrogen, Victoria, Australia, #12183-016; DRG pairs and isolated urothelial cells) or the PureLink RNA Mini kit (Invitrogen, #12183018A; bladder mucosa and detrusor) with DNAse treatment (Life Technologies, #12185-010) according to the manufacturer’s instructions. A NanoDrop Lite spectrophotometer (Thermofisher Scientific) was used to determine RNA purity and quantity. Using SuperScript VILO Master Mix (Invitrogen, #11755250), RNA was reverse transcribed to cDNA as per the manufacturer’s instructions. cDNA was then stored at −20°C for QRT-PCR.

#### Quantitative reverse-transcription polymerase chain reaction (QRT-PCR)

QRT-PCR was performed using Taqman Gene Expression Master Mix (Applied Biosystems, Victoria, Australia, #4369016) with commercially available hydrolysis probes (TaqMan; Life Technologies, see Table 1 for details) and RNAse-free water (AMBION, Victoria, Australia, #AM9916). For each reaction, 10 μL of qPCR MasterMix, 1 μL of TaqMan primer assay, 4 μL of water, and 5 μL of cDNA (1:2 dilution in RNA-free H_2_O) from each sample was tested in duplicate for each target. Endogenous controls Actb (β-actin; DRG pairs) and Gapdh and Hprt (glyceraldehyde 3-phosphate dehydrogenase and hypoxanthine phosphoribosyltransferase 1; mucosal and detrusor tissue layers and isolated urothelial cells) were used as endogenous controls. Assays were run for 45 cycles on a 7500 Fast Real-Time PCR System (Applied Biosystems) machine, using 7500 Fast software, v2.0.6. mRNA quantities are expressed as 2^−ΔCt^ relative to reference gene *Actb* (DRG pairs) *or Gapdh x Hprt* (geometric mean; mucosa, detrusor and urothelial cells) (Grundy et al., 2018c; Grundy et al., 2018e; Castro et al., 2019).

**Table 1.** Primers used for QRT-PCR and RT-PCR receptor expression assays.

| **Gene Alias** | **Gene Target** | **Assay ID** |
| --- | --- | --- |
| β-actin (reference gene) | *Actb* | Mm00607939_s1 |
| HPRT (reference gene) | *Hprt* | Mm00446968_m1 |
| Gapdh (reference gene) | *Gapdh* | Mm99999915_g1 |
| Tubulin-3 (neuronal marker) | *Tubb3* | Mm00727586_s1 |
| Mas-related G-protein Receptor A3 | *MrgprA3* | Mm02620679_s1 |
| Mas-related G-protein Receptor C11 | *MrgprX1* | Mm02525847_g1 |
| Mas-related G-protein Receptor D | *MrgprD* | Mm01701850_s1 |
| Transient receptor potential vanilloid 1 | *Trpv1* | Mm01246300_m1 |
| Transient receptor potential ankyrin 1 | *Trpa1* | Mm01227437_m1 |
| Transient receptor potential vanilloid 4 | *Trpv4* | Mm00499025_m1 |

### Cell culture of bladder-innervating DRG neurons

Four days following retrograde tracing, mice were humanely euthanised via CO_2_ inhalation and lumbosacral (LS; L5-S1) dorsal root ganglia (DRG) were removed. DRG were digested in Hanks’ balanced salt solution (HBSS; pH 7.4; Life Technologies, #14170161) containing 2.5 mg/mL collagenase II (GIBCO, ThermoFisher Scientific, #17101015) and 4.5 mg/mL dispase (GIBCO, ThermoFisher Scientific, #17105041) at 37°C for 30 min. Following aspiration of the collagenase-dispase solution, DRG were incubated in HBSS containing collagenase (4.5 mg/mL) only for 10 min at 37°C. After subsequent washes in HBSS, trituration through fire-polished Pasteur pipettes of descending diameter in 600μL complete DMEM (Dulbecco’s Modified Eagle Media [DMEM; GIBCO, ThermoFisher Scientific, #11995065]; 10% foetal calf serum [Invitrogen, ThermoFisher Scientific, MA, USA]; 2mM L-glutamine [GIBCO, ThermoFisher Scientific, #25030081], 100μM MEM non-essential amino acids [GIBCO, ThermoFisher Scientific, #11140076], 100mg/mL penicillin/streptomycin [GIBCO, ThermoFisher Scientific, #15070063], and 96 μg/L nerve growth factor-7S [Sigma, N0513-0.1MG]) mechanically disrupted DRG and dissociated cells, which were then centrifuged for 1 min at 50 g (Osteen et al., 2016; Inserra et al., 2017; Grundy et al., 2018a; Grundy et al., 2018c). Supernatant was gently aspirated, and neurons resuspended in 360 μL complete DMEM and spot-plated (30 μL) onto laminin (20μg/mL; Sigma-Aldrich, #L2020) and poly-D-lysine (800 μg/mL; ThermoFisher Scientific) coated 13 mm coverslips. Coverslips were incubated for 2-3 hours at 37°C in 5% CO_2_ to allow adherence of neurons before flooding with 1.7 mL complete DMEM. Cultured neurons were then maintained in an incubator at 37°C in 5% CO_2_ for 18-48 hours for calcium imaging, patch clamp recordings, or cell picking for single-cell RT-PCR (Grundy et al., 2018c; Castro et al., 2019).

### Single-cell RT-PCR of individual bladder-innervating dorsal root ganglia neurons

Under continuous perfusion of sterile and RNA-/DNase-free PBS, single retrogradely traced bladder DRG neurons were identified using a fluorescence microscope and collected into the end of a fine glass capillary using a micromanipulator (Grundy et al., 2018c). The glass capillary containing the cell was then broken into a sterile Eppendorf tube containing 10 μL of lysis buffer with DNAse (TaqMan Gene Expression Cells-to-CT Kit; Invitrogen, #4399002). A bath control was also taken for each coverslip of cells and analysed concurrently with cells. After lysis and DNAse treatment, samples were immediately frozen on dry ice and stored at −80°C until cDNA synthesis was performed. RNA was reverse transcribed to cDNA using SuperScript™ IV VILO™ Master Mix with ezDNase™ Enzyme (Invitrogen, #11766500) as per the manufacturer’s instructions. cDNA was then stored at −20°C for real time PCR. Tubulin-3 expression was used as a neuronal marker and positive control. Expression of each target gene within a single cell was determined by looking at the log (ΔRn) curve against cycle number. Only amplification curves that exhibited an exponential phase that rose steadily and eventually plateaued within 50 cycles were counted.

### Calcium imaging of cultured DRG neurons

Cultured DRG neurons (18-48 hours) were loaded with 2.5 μM Fura-2-acetoxymethyl ester (Fura-2; Invitrogen, ThermoFisher Scientific, #F1221) in 0.01% pluronic F-127 (Invitrogen, ThermoFisher Scientific, #P3000MP) at 37°C for 30 min followed by a 10 min wash with HEPES buffer (10 mM HEPES sodium salt [4-(2-hydroxyethyl)piperazine-1-ethanesulfonic acid sodium salt; Sigma, #H7006-100G], 140 mM NaCl [Chem Supply, #SA046-3KG], 4 mM KCl [Chem Supply, #PA054-500G], 5 mM D-glucose anhydrous [Chem Supply, #GA018-500G], 2 mM CaCl_2_ [Scharlau, #CA01951000], and 2 mM MgCl_2_ [Sigma, #M8266-100G], pH 7.40) before imaging at room temperature (23°C) (Bellono et al., 2017; Grundy et al., 2018c; Grundy et al., 2020). Emissions for Fura-2 were measured at 510 nm, following excitation at 340 and 380 nm, using a Nikon TE300 Eclipse microscope equipped with a Sutter DG-4/OF wavelength switcher, an Omega XF04 filter set for Fura-2, a Photonic Science ISIS-3 intensified CCD camera, and Universal Interface Card MetaFluor software. Retrogradely traced bladder DRG neurons were identified by the presence of the CTB-488, visible with excitation at 480 nm. Excitation at 480 nm was used only as an initial bladder neuron identifier; to prevent unnecessary photobleaching, images were not obtained at 480 nm throughout the experiment. Fluorescence images were obtained every 2 seconds using a 20x objective. Data were recorded and further analysed using MetaFluor software.

After an initial baseline reading to ensure cell fluorescence was stable, indicating healthy cells, DRG neurons were stimulated with chloroquine (1 μM; Sigma-Aldrich, #PHR1258-1G), BAM8-22 (2 μM; Tocris, #1763) and NPFF (5 μM; Tocris, #3137) with changes in intracellular calcium (Ca^2+^_i_) were monitored in real time. The TRPV1 receptor agonist capsaicin (3 μM; Sigma-Aldrich, M2028) was applied at the end of experiments. Ca^2+^_i_ is expressed as the ratio between the fluorescence signals at 340- and 380-nm (Fura-2 (340/380)).

### Patch-clamp electrophysiology of bladder-innervating dorsal root ganglia neurons

#### Current-clamp recordings

Whole cell patch-clamp recordings in current clamp mode were made from retrogradely traced fluorescently labelled lumbosacral (L5-S1) bladder-innervating DRG neurons using borosilicate glass pipettes (WPI) fire-polished to 3 to 7 MΩ and back-filled with an intracellular solution consisting of (in mM): 135 KCl; 2 MgCl2; 2 MgATP; 5 EGTANa; 10 HEPES-Na; adjusted to pH 7.3, 275 mOsm. Rheobase (amount of current required to fire an action potential) was determined in normal extracellular bath solution ((in mM): 140 NaCl; 4 KCl; 2 MgCl2; 2 CaCl2; 10 HEPES; 5 glucose, adjusted to pH 7.4, 285 mOsm) by application of a series of depolarising pulses from a holding potential of −65 mV (10 or 25 pA increments [480ms]) followed by hold at −65 mV (100 ms). Cells were continually perfused with bath solution by a gravity driven multi-barrel perfusion system positioned within 1 mm of the neuron under investigation (Osteen et al., 2016; Grundy et al., 2018c; Grundy et al., 2018e). After baseline recordings in external bath solution, an Mrgpr agonist ‘itch cocktail’ (MrgprA3 agonist; Chloroquine 10μM; Sigma-Aldrich, #PHR1258-1G; MrgprC11 agonist; BAM8-22: 2μM, Tocris, #1763; MrgprC11/A4 agonist; Neuropeptide FF (NPFF) 5 μM; Tocris, #3137) was applied for a 2 minute incubation period and a second recording was made.

### *In vivo* spinal dorsal horn activation

#### *In vivo* bladder infusion

*In vivo* bladder infusion in anaesthetised mice was performed as previously described (Grundy et al., 2018c; Grundy et al., 2020). 12-week old female mice were anaesthetized (isoflurane 2% - 4% in oxygen) and a catheter (AS point style, 51 mm, metal hub, 21-gauge needle [Hamilton, 7751-12]) was inserted into the bladder via the urethra. Correct catheter placement was determined by inserting gently until meeting resistance (bladder dome) and receding slightly (2-3 mm) to avoid damaging the bladder wall. Urine was removed using a suction syringe. Subsequently, a new catheter (AS point style, 51 mm, metal hub, 21-gauge needle) was inserted, primed with either vehicle (saline) or an Mrgpr agonist cocktail (BAM8-22: 20μM; NPFF: 50μM; and Chloroquine: 100μM), and 100 μL of solution was then gently infused, without fully distending the bladder, and allowed to incubate for 5 minutes. Immediately following removal of the compound, mice were administered an anaesthetic overdose (intraperitoneal injection of pentobarbitone, 1.1 mg/g body weight; Lethabarb®; Virbac Australia) followed by 4% buffered paraformaldehyde transcardial perfuse fixation.

##### Transcardial perfuse fixation and tissue dissection

After anaesthetic overdose, the thoracic cavity was opened and 0.5 mL heparinised saline (50 IU in 5 mL; Pfizer) was injected into the left ventricle. A 22-gauge needle attached to tubing and a peristaltic perfusion pump was then inserted into the injection site. The right atrium was cut open, allowing for perfusate drainage. Warm phosphate buffer (0.1 M; 21.7 mM NaH_2_PO_4_ [Chem Supply, #SA061-500G], 81 mM Na_2_HPO_4_ [VWR Chemicals, #102494C], pH 7.2) was perfused, followed by ice-cold 4% paraformaldehyde (PFA; Sigma-Aldrich, #158127) in 0.1 M phosphate buffer. Following complete PFA perfusion, the lumbosacral spinal cord (determined by the level of dorsal root ganglia (DRG) root insertion points; the lowest rib was used as an anatomical marker of DRG level T13) was removed and post-fixed in 4% PFA in 0.1 M phosphate buffer at 4°C for 18 to 20 hours. After post-fixation, spinal cord segments were cryoprotected in 30% sucrose/phosphate buffer (Sigma-Aldrich, #S9378) for 48 hours at 4°C, followed by a 48-hour incubation at 4°C in 50% optimal cutting temperature (OCT; VWR, #C361603E)/30% sucrose/phosphate buffer solution prior to freezing in 100% OCT using liquid nitrogen-cooled 2-methylbutane (Sigma-Aldrich, #M32631). Frozen spinal cord segments were cryosectioned (10 μm thick) and placed onto gelatin-coated slides for immunostaining. Sections were performed serially and distributed over 6 slides, which were used for immunohistochemical localisation of phosphorylated-MAP-kinase ERK 1/2 (pERK) (Grundy et al., 2018c; Grundy et al., 2019b; Harrington et al., 2019; Grundy et al., 2020). The number of pERK-immunoreactive neurons were counted in L6-S2 spinal cord sections, with spinal segments determined *ex vivo* by the shape of the dorsal horn (Lein et al., 2007).

##### Immunohistochemistry of phosphorylated-MAP-kinase ERK 1/2 within the spinal cord

The dorsal horn neurons activated by bladder infusion were identified by labelling for neuronal activation marker, phosphorylated-MAP-kinase ERK 1/2 (pERK), using the DAKO Omnis autostainer (Agilent Technologies Australia, Mulgrave, Vic, AUS), with 3,3′-Diaminobenzidine (DAB) / horseradish peroxidase (HRP) secondary antibody staining. Sections were processed via the University of Adelaide Health and Medical Sciences Histology services. pERK activation was determined rather than c-Fos due to the rapidity of ERK phosphorylation (minutes compared to hours) (Gao and Ji, 2009). Given the short-duration nature of the specific peripheral stimuli applied, the rapid onset of pERK provides a more appropriate correlate. Further, quantification of pERK activation is increasingly being utilised to identify spinal activation to a range of peripheral stimuli (Jiang et al., 2015; Castro et al., 2017; Grundy et al., 2018a; Grundy et al., 2018c; Grundy et al., 2020). After air drying for 1 hour, sections were post-fixed in formalin with a brief rinse in distilled water (dH2O) prior to target retrieval at 97°C for 30 minutes with EnVision FLEX TRS Low pH target retrieval solution (citrate buffer, pH 6.1; K8005, Agilent DAKO), followed by a wash step with wash buffer (GC807, DAKO Omnis, Agilent). A 10-minute protein block step with Serum-Free Protein Block (X0909, Agilent DAKO) to prevent non-specific binding was followed by a 1-hour incubation in primary antibody (pERK; 1:800) in Antibody Diluent (S0809, Agilent DAKO). Following removal of excess primary antibody with wash buffer, a 3-minute incubation in Envision FLEX Peroxidase-blocking Reagent (GV823, Agilent DAKO) was performed. Slides were then washed twice with wash buffer before a 20-minute incubation with Envision FLEX HRP Polymer (GV823, Agilent DAKO) for HRP binding. A further 2 wash steps were performed, followed by a DI rinse and a third wash step, before a 10-minute incubation in EnVision FLEX Substrate Working Solution (DAB) for staining. Slides underwent alternate cycles of wash buffer, DI rinse and wash buffer prior to a 3-minute incubation with Haematoxylin ready-to-use solution (K8018, Agilent DAKO) and a further DI rinse and wash step. Slides were removed from the DAKO Omnis, dehydrated in alcohol, and cleared in xylene before mounting with dibutylphthalate polystyrene xylene (DPX, Sigma-Aldrich) and cover slipping.

##### Microscopy

DAB/HRP-stained slides were imaged using a NanoZoomer microscope (NanoZoomer 2.0-HT, Hamamatsu), with 1 – 4 manually set focus points per section. Images of individual sections from L6 – S2 regions were taken using NanoZoomer digital pathology viewing software (NDP.view2, Hamamatsu) and analysed using ImageJ software. The images were not manipulated in any way.

#### Spinal cord pERK neuronal counts and analysis

Neuronal counts were analysed from previously saved digital photomicrographs, with only neurons with intact nuclei counted. The number of pERK-immunoreactive (pERK-IR) neurons in a quadrant of the L6-S2 dorsal horn was obtained from minimum 6 sections/animal viewed at 10x magnification. The mean number of pERK-IR neurons (± SEM) in the superficial dorsal horn (SDH; LI-II), the dorsal grey commissure (DGC), and the sacral parasympathetic nuclei (SPN) in sacral spinal segments were compared between vehicle- and Mrgpr agonist cocktail-treated mice. These regions are known to have roles in nociceptive signalling and autonomic reflexes (Todd, 2010) and are known to be activated by bladder afferent output (Grundy et al., 2018a; Grundy et al., 2019a).

### Experimental design and statistical analyses

Data are presented as Mean ± SEM or the % of afferents or neurons. Within the specific figure legends, N indicates the number of animals, whilst n indicates the number of independent afferents or neurons. Sample size was based on historical data and the use of power calculations. Statistical significance was reported at levels of \**P* <0.05, \*\**P* <0.01, \*\*\**P* <0.001, \*\*\*\**P* <0.0001. In some cases, due to space limitation, # is used to indicate significance of *P*<0.0001 for multiple comparisons. Data were tested for Gaussian distribution using Prism 8 (GraphPad, San Diego, CA, USA) in order to determine if they were normally distributed or not and therefore the correct statistical tests to be used. Data were then analysed accordingly using Prism 8 (GraphPad, San Diego, CA, USA), via one- or two-way analysis of variance (ANOVA) with Tukey’s or Sidak’s post hoc analyses dependent on data distribution, or student’s t-tests, for parametric data. Non-parametric data were analysed using Kruskal-Wallis test with Dunn’s multiple comparisons, Mann-Whitney analysis, or Wilcoxon matched-pairs signed rank test. The specific tests used for analysis of each data set is indicated within the individual figure legends.

## RESULTS

### Mrgpr agonists induce distension evoked mechanical hypersensitivity of bladder afferents

In order to determine if pruritogenic receptors have a functional role in bladder sensory signalling, we performed *ex vivo* bladder afferent recordings and determined their mechanosensitivity to bladder distension before and during bladder instillation with Mrgpr agonists. Intravesical application of the MrgprA3 agonist, chloroquine, evoked bladder afferent hypersensitivity to bladder distension in whole nerve recordings as determined by the total area under the curve of the response **(Figure 1A, 1B)**. Analysis of the pressure/volume relationship in the bladder during repetitive graded bladder distensions before and during chloroquine revealed no effect on bladder compliance **(Figure 1C)**. This indicates that the mechanical hypersensitivity induced by bladder infusion of chloroquine is not secondary to changes in bladder muscle function, which can indirectly influence bladder afferent responsiveness. To further investigate the afferent response, we performed post-hoc single unit analysis, which revealed that distinct populations of bladder afferents became hypersensitive following chloroquine application (called responders; **Figure 1Di, 1Dii)**, whilst other afferents were unaffected (called non-responders; **Figure 1Ei, 1Eii)**. Of the afferents that were affected by chloroquine the predominant effects were apparent throughout most distension pressures from 14-30mmHg **(Figure 1Di, 1F)**. Overall, 36% of bladder afferents were responsive to chloroquine application **(Figure 1G)**.

**Figure 1:**
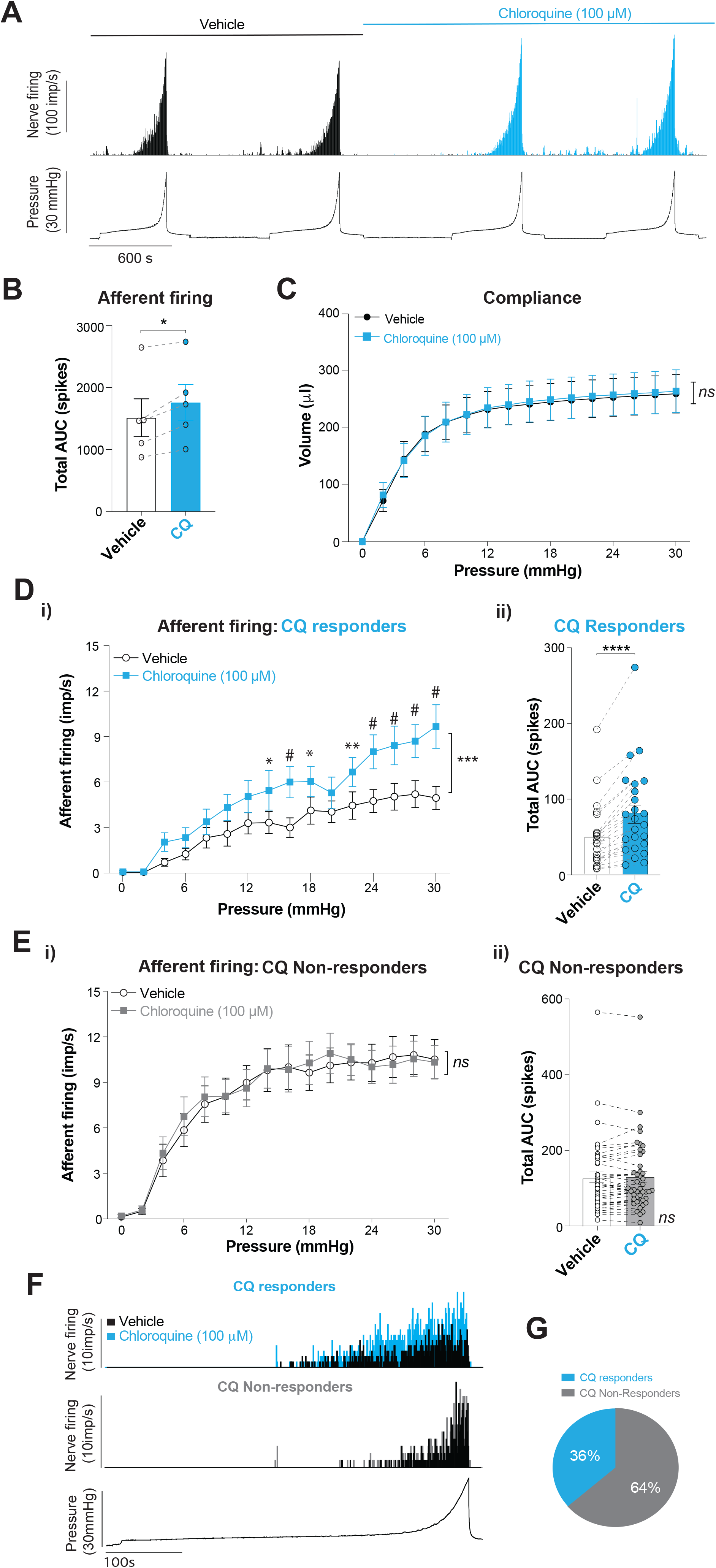
The MRGPRA3 agonist chloroquine (CQ) evokes mechanical hypersensitivity in a cohort of bladder afferents. **A)** Original recordings from an *ex vivo* nerve recording preparation showing bladder afferent firing and intravesical pressure. Recordings show exaggerated whole nerve bladder afferent responses to distension following intravesical infusion with chloroquine (100μM). **B)** The sum total of afferent firing (Total Area Under Curve; 0-30mmHg) for each individual experiment showing bladder instillation with chloroquine (100μM) significantly increased bladder afferent firing to distension (\**P*<0.05, paired t-test, N=5 mice). Dots indicate whole nerve responses from individual mice before and after the itch cocktail. **C)** The relationship between intra-bladder pressure and volume (compliance) during graded bladder distension (30μL/min from 0-30mmHg) is unaffected by intravesical administration of chloroquine (100μM, *ns, P*>0.05, repeated-measures 2-way ANOVA, N=5). **D) i)** Post-hoc single unit analysis of the multiunit *ex vivo* bladder recordings shown in panel A identified bladder afferents that were responsive to chloroquine (showing a >20% increase in afferent firing) in the presence of chloroquine (\*\*\**P*<0.001, two-way ANOVA with \**P*<0.05, \*\**P*<0.01, ^#^*P*<0.0001, Sidak’s multiple comparisons test, n=24 afferents, N=5 mice). **ii)** Chloroquine responsive bladder afferents showing enhanced firing as determined by the total number of action potentials fired (\*\*\*\**P*<0.0001, Wilcoxon matched-pairs signed rank test, n=24 afferents, N=5 mice). **E)** Post-hoc single unit analysis also identified bladder afferents unaffected by chloroquine when analysed **i)** across the entire distension range (*ns*, *P*>0.05, Two-way RM ANOVA, n=42 afferents, N=5 mice) or as **ii)** the total number of action potentials fired (*ns*, *P*>0.05, Wilcoxon matched-pairs signed rank test, n=42 afferents, N=5 mice). **F)** Original recordings of chloroquine (100μM) responding (upper panel) and non-responding (middle panel) bladder afferents during controlled bladder distension from 0-30-mmHg (lower panel). **G)** Overall, 36% of single units (24 of 66) bladder afferents showed a >20% increase in afferent firing to chloroquine. Data represent Mean ± SEM. Chloroquine: CQ. Area Under Curve: AUC. Impulses per second: imp/s^−1^.

To follow up on these findings we also used the MrgprC11 agonist BAM8-22 and found that it also significantly enhanced bladder afferent responses to bladder distension **(Figure 2A, 2B)**, without affecting muscle compliance **(Figure 2C)**. Single unit analysis of bladder afferents revealed that BAM8-22 also sensitized a sub-population of bladder afferents **(Figure 2Di, 2Dii)**, whilst another population was unaffected **(Figure 2Ei, 2Eii)**. Responsive afferents were particularly affected at distension pressures at or above 18mmHg **(Figure 2Di, 2F)** and constituted 20% of the bladder afferents recorded **(Figure 2G)**. We also found that the MrgprC11/MrgprA4 agonist neuropeptide FF (NPFF) evoked significant bladder afferent hypersensitivity **(Figure 3A, 3B)**, that was independent of changes in muscle compliance **(Figure 3C)** and affected the responsiveness of 41% of bladder afferents recorded **(Figure 3D, 3E, 3F, 3G)**. Neither chloroquine, BAM8-22, nor NPFF recruited ‘silent’ afferents to become subsequently mechanosensitive. Overall, these findings show that chloroquine, BAM8-22 and NPFF all evoke hypersensitivity to distension in sub-populations of bladder afferent, suggesting the Mrgpr receptors they activate are expressed on specific subpopulations of bladder-innervating DRG neurons.

**Figure 2:**
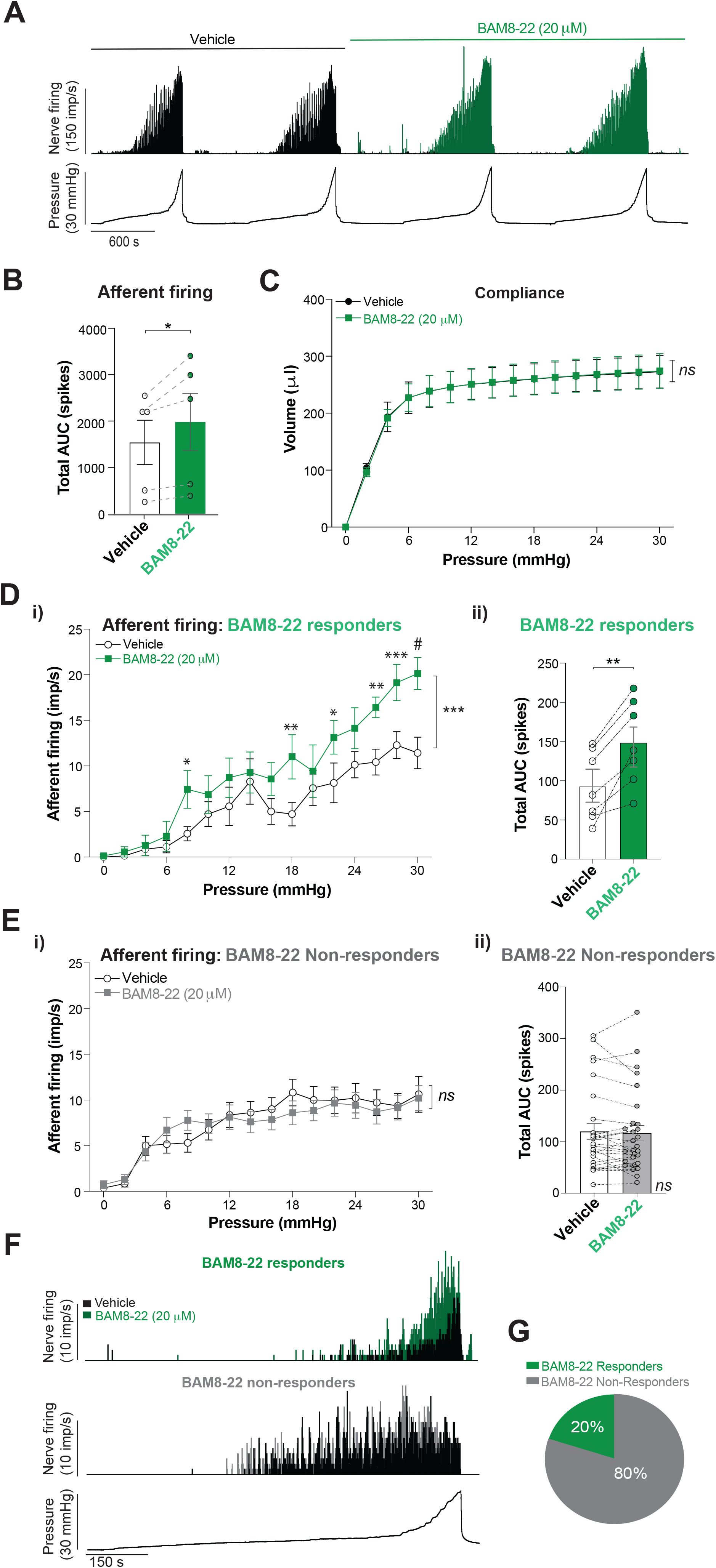
The MRGPRC11 agonist BAM8-22 evokes mechanical hypersensitivity in a cohort of bladder afferents. **A)** Original *ex vivo* recordings showing exaggerated bladder afferent responses to distension following intravesical infusion with BAM8-22 (20μM). **B)** Total bladder afferent firing was significantly increased following bladder instillation with BAM8-22 (20μM, \**P*<0.05, paired t-test, N=5 mice). **C)** Bladder compliance during graded bladder distension is unaffected by intravesical administration of BAM8-22 (20μM, *ns, P*>0.05, repeated-measures 2-way ANOVA, N=5). **D)** Post-hoc single unit analysis demonstrated a population of BAM8-22 responsive bladder afferents showing enhanced firing as determined **i)** across the distension range (\*\*\**P*<0.001, Two-way RM ANOVA, with \**P*<0.05, \*\**P*<0.01, \*\*\**P*<0.001, ^#^*P*<0.0001 based on Sidak’s multiple comparisons test n=7 afferents, N=5 mice) or as **ii)** the total number of action potentials fired (\*\**P*<0.01, Paired t-test, n=7 afferents, N=5 mice). **E)** Post-hoc single unit analysis also identified bladder afferents unaffected by BAM8-22 when analysed **i)** across the entire distension range (*ns*, *P*>0.05, Two-way RM ANOVA, n=28 afferents, N=5 mice) or as **ii)** the total number of action potentials fired (*ns*, *P*>0.05, Wilcoxon matched-pairs signed rank test, n=28 afferents, N=5 mice). **F)** Original recordings of BAM8-22 responding and non-responding bladder afferents during bladder distension from 0-30 mmHg. **G)** Overall, 20% (7 of 35) of bladder afferents showed a >20% increase in afferent firing to BAM8-22. Data represent Mean ± SEM. Impulses per second: imp/s^−1^.

**Figure 3:**
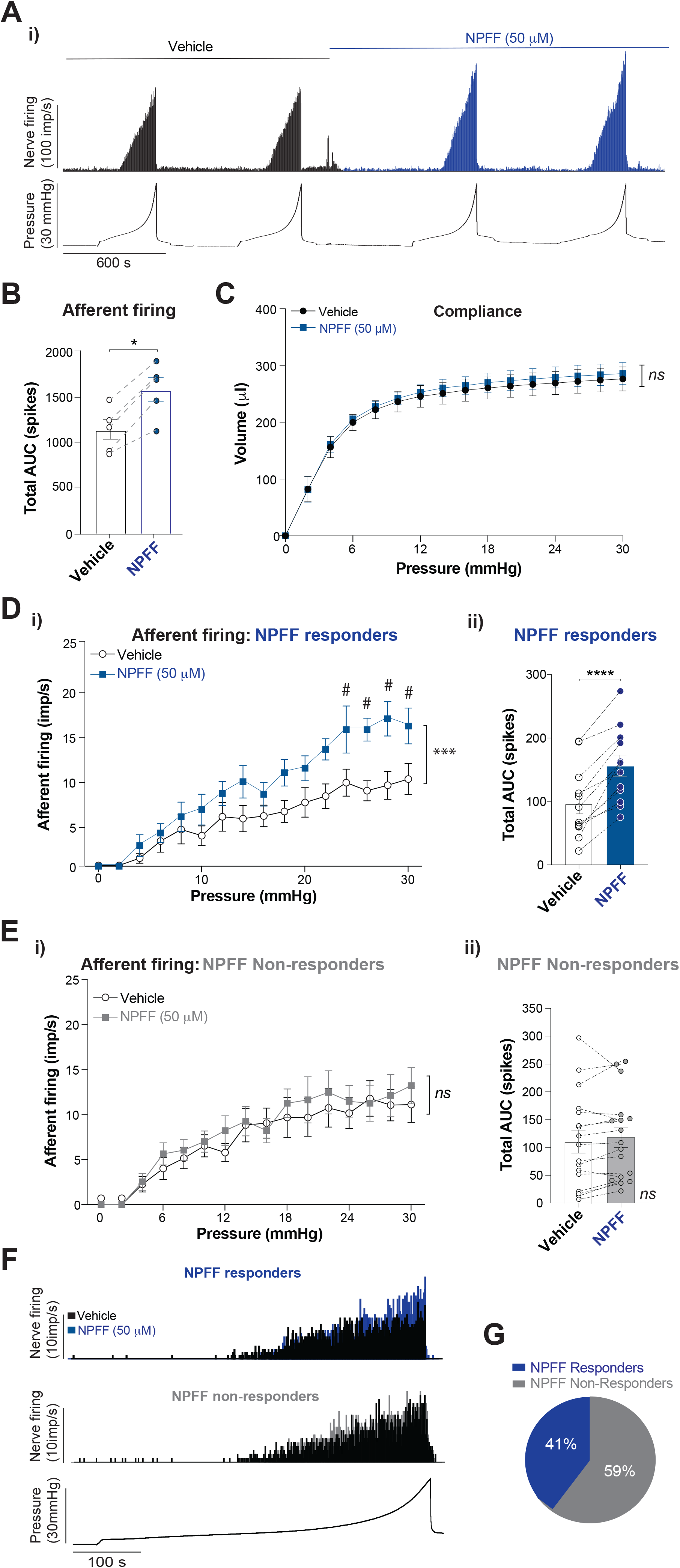
The MRGPRC11/A4 agonist NPFF evokes mechanical hypersensitivity in a cohort of bladder afferents. **A) i)** *Ex vivo* recordings showing an enhanced afferent response to distension following intravesical infusion with NPFF (50μM). **B)** Total bladder afferent firing was significantly increased following bladder instillation with NPFF (50μM, \**P*<0.05, paired t-test, N=5 mice). **C)** Bladder compliance is unaffected by intravesical administration NPFF (5μM, *ns, P*>0.05, repeated-measures 2-way ANOVA, N=5 for each group). **D)** Post-hoc single unit analysis showed a population of NPFF responsive bladder afferents showing enhanced firing **i)** across distension pressures (\*\*\**P*<0.001, Two-way RM ANOVA, with ^#^*P*<0.0001 based on Sidak’s multiple comparisons test n=12 afferents, N=5 mice) or as **ii)** the total number of action potentials fired (\*\*\*\**P*<0.0001, Paired t-test, n=12 afferents, N=5 mice). **E)** A population of bladder afferents were unresponsive to NPFF based on **i)**responses across the distension range (ns, *P*>0.05, Two-way RM ANOVA, n=17 afferents, N=5) or **ii)**the total number of action potentials fired (ns, *P*>0.05, Paired t-test, n=17 afferents, N=5 mice). **F)** Example traces of responding and non-responding bladder afferents to NPFF (50μM) during bladder distension. **G)** Overall, 41% of single units (12 of 29) bladder afferents showed a >20% increase in afferent firing to NPFF. Data represent Mean ± SEM. Impulses per second: imp/s^−1^.

In order to confirm that these effects were mediated via Mrgpr activation we performed additional *ex vivo* bladder afferent recordings with tissue from *Mrgpr-clusterΔ*^*−/−*^ mice **(Figure 4)**. In our previous studies focusing on colonic sensitivity, we have shown that a combined application of Mrgpr agonists, as an ‘itch cocktail’, causes pronounced colonic mechanical hypersensitivity, an effect which is absent in *Mrgpr-clusterΔ*^*−/−*^ mice (Castro et al., 2019). In the current study we found that *ex vivo* bladder afferent responses to graded distension in *Mrgpr-clusterΔ*^*−/−*^ mice were identical when comparing between the application of vehicle or the ‘itch cocktail’ of co-applied chloroquine, BAM8-22 and NPFF **(Figure 4A, 4Bi, Bii)**. Additionally, these studies also showed that bladder compliance was unaltered in vehicle or ‘itch cocktail’ treated bladders from *Mrgpr-clusterΔ*^*−/−*^ mice **(Figure 4C)**. These findings confirm the role of Mrgprs in chloroquine, BAM8-22 and NPFF induced bladder afferent mechanical hypersensitivity.

**Figure 4:**
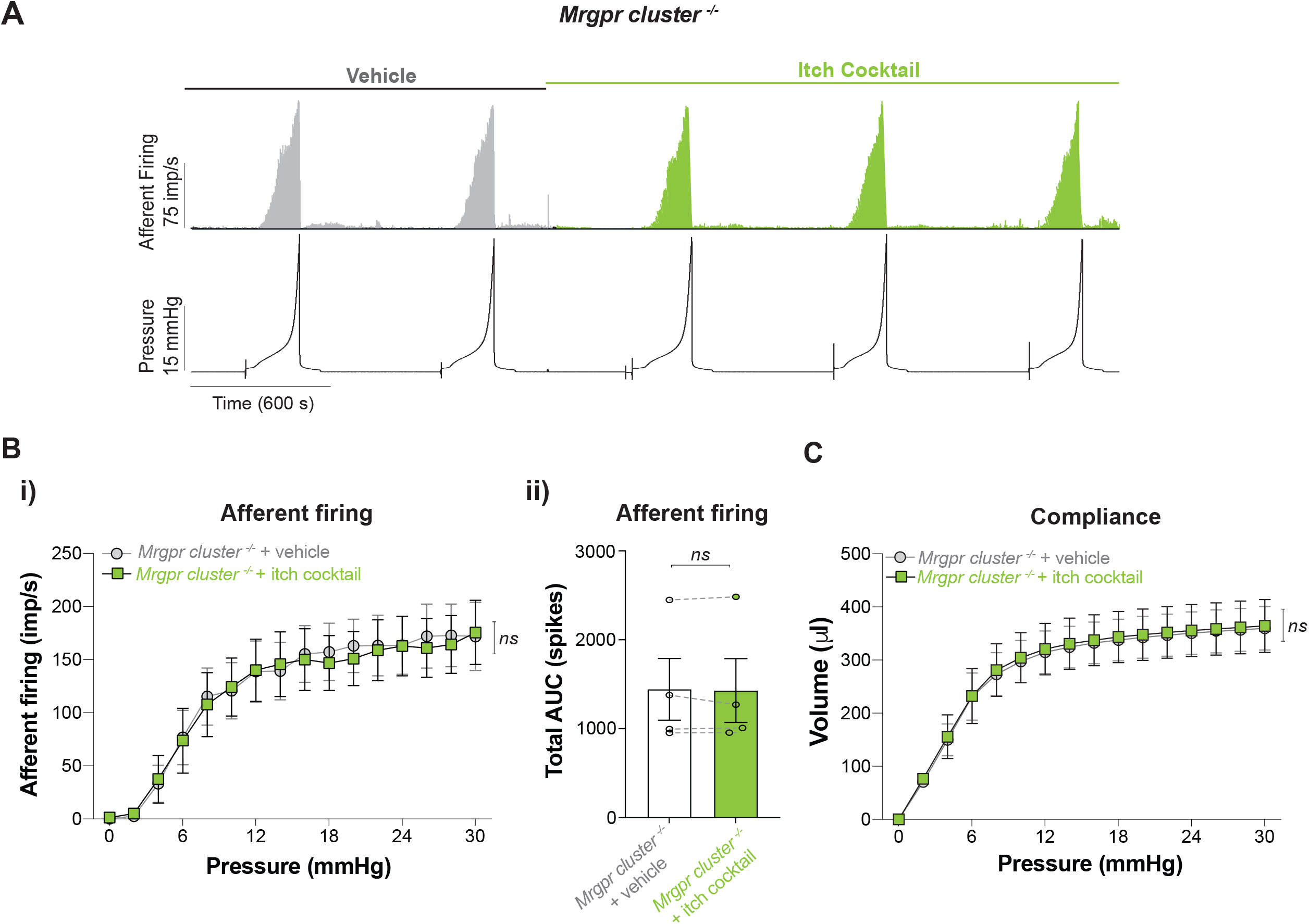
Bladder afferent responses to an ‘itch cocktail’ are absent in *Mrgpr-clusterΔ*^*−/−*^ mice. **A)** Original recordings of bladder afferent firing and intravesical pressure during consecutive bladder distensions (0-30mmHg) with saline followed by the ‘itch cocktail’ containing the Mrgpr agonists chloroquine (100μM0, BAM8-22 (20μM) and NPFF (50μM). Afferent responses were stable during saline infusion and unchanged after administration of the Mrgpr agonists. **B) i)** Group data from *ex vivo* afferent recordings using *Mrgpr-clusterΔ*^*−/−*^ mice showing almost identical responses to bladder instillation of vehicle (saline) or an ‘itch cocktail’ (ns, *P*>0.05, 2-way RM ANOVA, N=4 mice). **ii)** Total bladder afferent firing was unchanged following bladder instillation of an ‘itch cocktail’ (ns, *P*>0.05, paired t-test, N=4 mice). Dots indicate whole nerve responses from individual mice before and after the itch cocktail. **C)** Bladder compliance during graded bladder distension (30μL/min, 0-30mmHg) was unaffected by intravesical administration of the Mrgpr agonist itch cocktail in *Mrgpr-clusterΔ*^*−/−*^ mice (*ns*, *P*>0.05, 2-way RM ANOVA, N=4). Data represent Mean ± SEM.

### MrgprA3 and MrgprC11 are expressed in sub-populations of bladder-innervating DRG neurons

To determine the mechanisms by which Mrgpr agonists induce distension-evoked bladder afferent hypersensitivity, we determined Mrgpr expression in DRG, as well as in bladder tissue. Lumbosacral (LS) DRG contain the cell bodies of sensory neurons innervating a number of different pelvic organs, including >30% of which innervate the mouse bladder (Christianson et al., 2007; Grundy et al., 2018e). Therefore, we performed retrograde tracing from the bladder wall in order to isolate those that innervate the bladder. Single-cell RT-PCR from retrogradely traced bladder-innervating DRG neurons revealed that of the 77 individual neurons studied, 14% expressed *MrgprA3*, whilst 38% expressed *MrgprC11* **(Figure 5A, 5B)**. In comparison mRNA transcripts for *Trpv1 and Trpa1*, key channels involved in bladder afferent function (Daly et al., 2007; DeBerry et al., 2014), were detected in 84% and 15% of bladder-innervating DRG neurons respectively. Interestingly, another pruritogenic Mrgpr, *MrgprD*, was only expressed in 6% of bladder-innervating DRG neurons, so was not investigated further with functional studies **(Figure 5A, 5B)**. Co-expression analysis revealed that the majority of bladder-innervating neurons that express *MrgprA3* co-express *MrgprC11* (90%) and *Trpv1* (81%), with a smaller proportion co-expressing *Trpa1* (36%) or *MrgprD* (18%) **(Figure 5B, 5C)**. Bladder-innervating *MrgprC11* expressing DRG neurons commonly co-expressed with *Trpv1* (89%), but less so with *MrgprA3* (34%), *TrpA1* (25%) or *MrgprD* (7%) **(Figure 5B, 5D)**. Of the few cells that expressed *MrgprD*, 40% co-expressed with either *MrgprA3*, *MrgprC11*, *TrpV1* or *TrpA1* **(Figure 5B, 5E)**. Overall, no bladder-innervating DRG neuron expressed the full repertoire of receptors investigated. Interestingly, the Mrgpr expression profile in bladder-innervating DRG neurons contrasts starkly with that observed in the whole LS DRG, whereby *MrgprD* mRNA is expressed in the highest abundance **(Figure 5F)**. In terms of bladder tissue, *MrgprA3*, *MrgprC11* and *MrgprD* were either expressed below the detection limit or were poorly expressed in the detrusor **(Figure 5G)**, the mucosa **(Figure 5H)**, or primary urothelial cells **(Figure 5I)**, particularly when compared with an abundantly expressed target such a *Trpv4* (Everaerts et al., 2010a; Everaerts et al., 2010b). Mrgpr mRNA expression was approximately 1000-100,000x lower than expression of *Trpv4.* Overall, these findings indicate that the pruritogenic receptors MrgprC11 and MrgprA3 are expressed on sensory neurons innervating the bladder, and their expressions profile correlates well with our observations that subpopulations of bladder afferents display mechanical hypersensitivity following application of the respective Mrgpr agonists. The relatively low expression of Mrgprs in bladder tissue correlates well with our findings that Mrgpr agonists do not affect bladder compliance.

**Figure 5:**
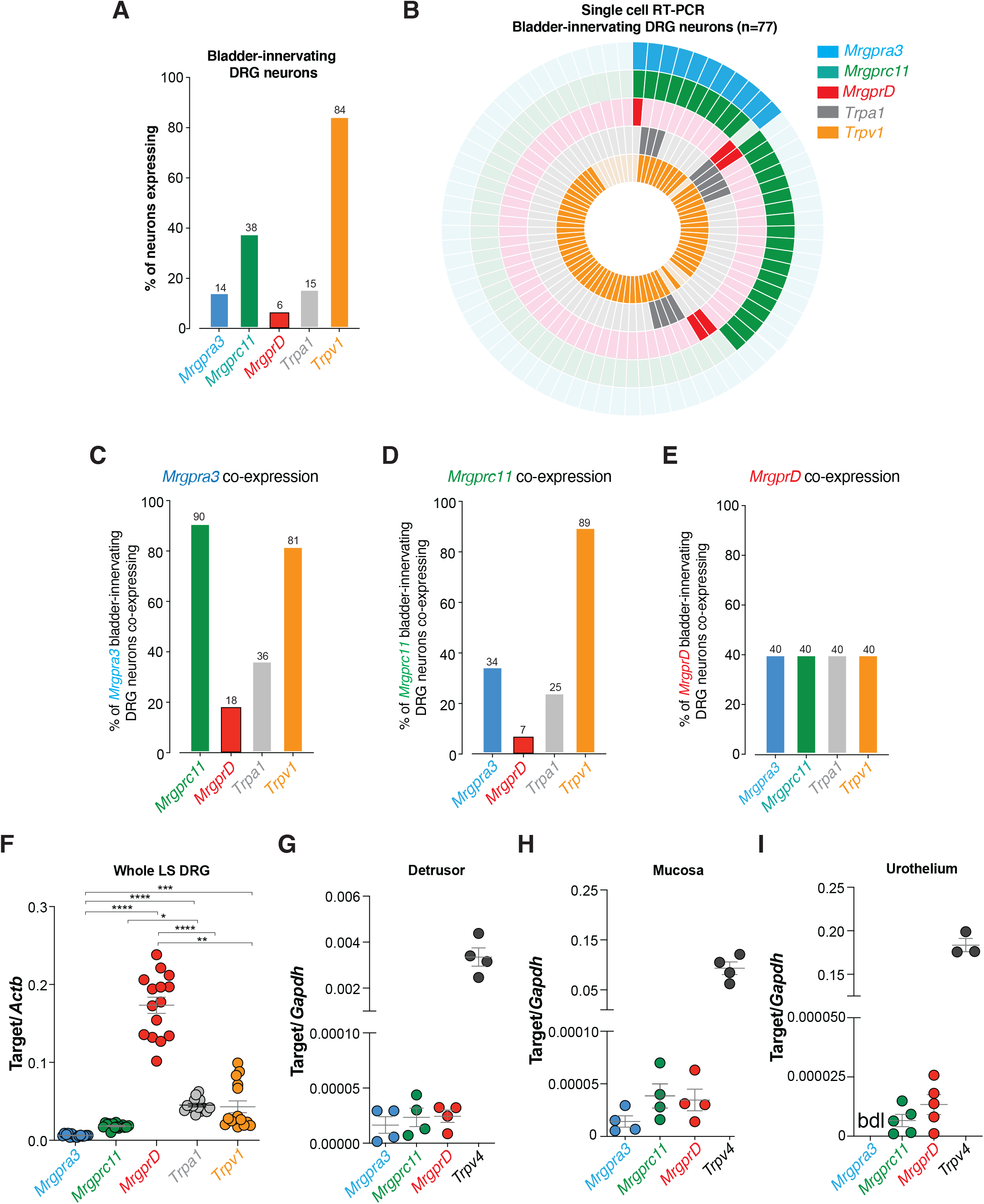
Mrgpr receptors are expressed by bladder-innervating DRG neurons. **A)** Single-cell RT-PCR of retrogradely traced bladder-innervating DRG neurons reveals the percentage of neurons expressing and co-expressing genes encoding *Mrgprs* and *Trp* channels (n=77 single cells from N=6 mice). mRNA for *MrgprA3* and *MrgprC11* were expressed in 14% and 38% of bladder-innervating DRG neurons respectively. In contrast, *MrgprD* was expressed in only 6% of bladder-innervating DRG neurons. *Trpa1* and *Trpv1* mRNA transcripts were detected in 15% and 84% of bladder-innervating DRG neurons. **B)** Donut plot showing expression and co-expression of genes encoding *MrgprC11, MrgprA3, MrgprD, Trpa1, and Trpv1* in 77 individual retrogradely traced bladder-innervating DRG neurons. Each colour represents an individual gene with expression marked by bold colouring. *MrgprA3* is represented in the outer ring with *Trpv1* in the inner ring. Individual neurons are arranged radially, such that co-expression of genes in a single neuron can be easily identified running from outside to inside. **C)** The majority of bladder-innervating DRG neurons that express *MrgprA3* co-express *MrgprC11* (90%) and *Trpv1* (81%). In contrast, a minor proportion of *MrgprA3* expressing bladder DRG neurons also co-express *MrgprD* (18%) or *Trpa1* (36%). **D)** *MrgprC11* is commonly co-expressed with *Trpv1* (89%), but less so with *MrgprA3* (34%), *MrgprD* (7%), and *Trpa1* (25%). **E)** Of the very few cells that express *MrgprD* (5 of 77 neurons), 40% (2 of 5 neurons) co-express with either *MrgprA3*, *MrgprC11*, *Trpv1* or *Trpa1*. **F)** mRNA expression of *MrgprA3*, *MrgprC11*, *MrgprD*, *Trpv1*, and *Trpa1* in whole LS (L5-S1) DRG. Expression of *MrgprD* mRNA is significantly higher than *MrgprC11*, *MrgprA3*, *Trpv1* and *Trpa1*. *MrgprC11* and *MrgprA3* are significantly less abundant than *Trpv1* and *Trpa1* (\**P*<0.05, \*\**P*<0.01, \*\*\**P*<0.001, \*\*\*\**P*<0.0001, Kruskal-Wallis test with Dunn’s multiple comparisons post hoc test, n=15 pairs of DRG from N=5 mice). **G-I)** mRNA expression of *MrgprA3, MrgprC11, MrgprD* (and the abundantly expressed *Trpv4* as a comparator) in bladder **(G)** detrusor smooth muscle, **(H)** mucosa and **(I)** primary urothelial cells. Expression of all Mrgprs were similar between the detrusor, urothelial cells and mucosa (tissue samples from N=3-5 mice). All data represent Mean ± SEM.

### Mrgpr agonists activate sub-populations of bladder-innervating DRG neurons

In order to confirm the results of our single-cell RT-PCR at a functional level, and to investigate the roles of Mrgpr agonists in activating bladder-innervating DRG neurons, we measured intracellular calcium ([Ca^2+]^i) using Fura-2AM in response to application of chloroquine, BAM8-22 and NPFF. Overall, calcium imaging of dissociated bladder-innervating DRG neurons revealed that individual application of chloroquine **(Figure 6A)**, BAM8-22 **(Figure 6B)** or NPFF **(Figure 6C)** resulted in activation of sub-populations of bladder-innervating DRG neurons. Overall, 8.1%, 22.9% and 16.2% of bladder-innervating LS DRG neurons were activated by chloroquine, BAM8-22 and NPFF respectively **(Figure 6D, 6E)**. In series application of multiple Mrgpr agonists demonstrated functional co-expression of MrgprA3 and MrgprC11 in populations of bladder-innervating DRG neurons **(Figure 6E–5J)**, supporting our single cell RT-PCR data suggesting colocalization of MrgprC11 and MrgprA3 in bladder-innervating DRG neurons.

**Figure 6:**
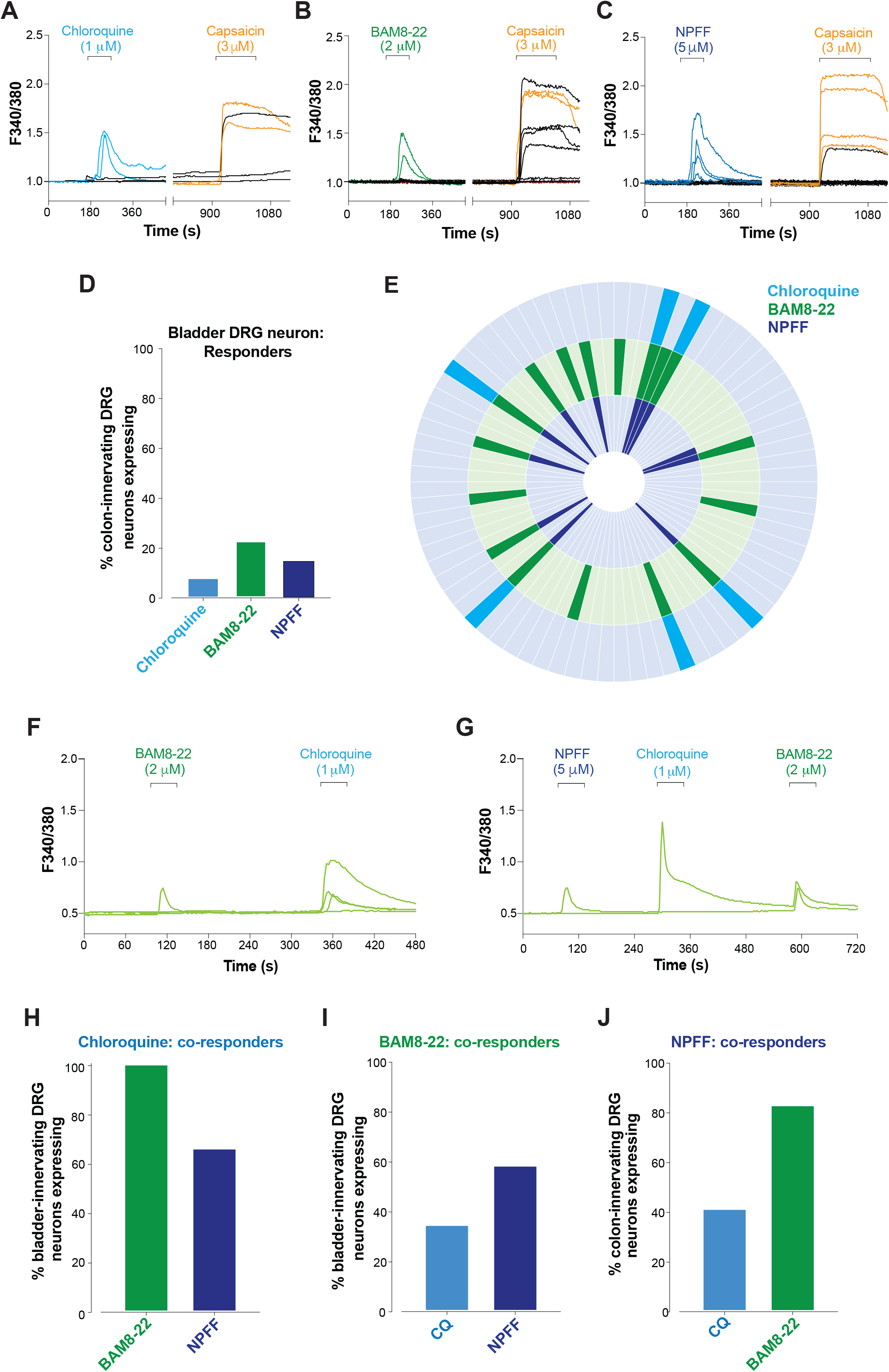
A cohort of bladder-innervating DRG neurons respond to MRGPR agonists. **A-C)** Calcium responses elicited by **(A)** chloroquine (CQ; 1μM) (**B)** BAM8-22 (2μM) and **(C)** NPFF (5μM), in isolated cultured retrogradely traced bladder-innervating LS DRG neurons loaded with Fura-2. Capsaicin was added at the end of each experiment to investigate the functional co-expression of TRPV1 with the MRGPR’s. Each line represents an individual neuron, coloured lines indicate bladder DRG neurons responding to MRGPR agonists. Orange capsaicin traces indicate bladder neurons that responded to either BAM8-22, chloroquine or NPFF. **D)** Percentage of bladder-innervating DRG neurons responding to chloroquine (8.1%: 6/74 neurons), BAM8-22 (22.9%: 17/74 neurons) or NPFF (16.2%: 12/74 neurons). **E)** Donut plot showing functional responsiveness of individual neurons to chloroquine, BAM8-22 and NPFF. Bold colouring represents a functional response to an Mrgpr agonist. Chloroquine is represented in the outer ring with NPFF in the inner ring. Individual neurons are arranged radially, such that functional responsiveness of a single neuron to the Mrgpr agonists can be easily identified running from outside to inside. **F-G)** Example traces showing bladder-innervating DRG neurons can respond to more than one MRGPR agonist. **H)** All bladder-innervating DRG neurons responding to chloroquine also responded to BAM8-22 (100%: 6/6 neurons), with 66.6% of chloroquine responding neurons also responding to NPFF (4/6 neurons). **I)** 35.2% of BAM8-22 responding neurons also responded to chloroquine (6/17 neurons), with 58.8% responding to NPFF (10/17 neurons). **J)** Of the NPFF responding neurons, 41.6% (5/12 neurons) also responded to chloroquine, with 83.3% (10/12 neurons) also responding to BAM8-22.

To further investigate the role of Mrgprs in modulating bladder sensation we used whole-cell patch-clamp electrophysiology of retrogradely traced bladder-innervating DRG neurons. In these studies, we co-applied the Mrgpr agonists chloroquine, BAM8-22 and NPFF at the same time as an ‘itch cocktail’, as used in some of our bladder afferent recording studies **(Figure 7A)**. We found that this cocktail caused significant hyperexcitability in a sub-population (44%) of bladder-innervating DRG neurons, as indicated by a significant decrease in the rheobase, or amount of current required to fire an action potential **(Figure 7Ai, 7Aii, 7Aiii)**. These changes were also accompanied by a significant increase in the number of action potentials fired at 2x rheobase **(Figure 7Bi, 7Bii, 7Biii)**. In comparison the neuronal excitability of another population of bladder-innervating DRG neurons were unaffected by the Mrgpr agonist cocktail **(Figure 7A, 7B)**.

**Figure 7:**
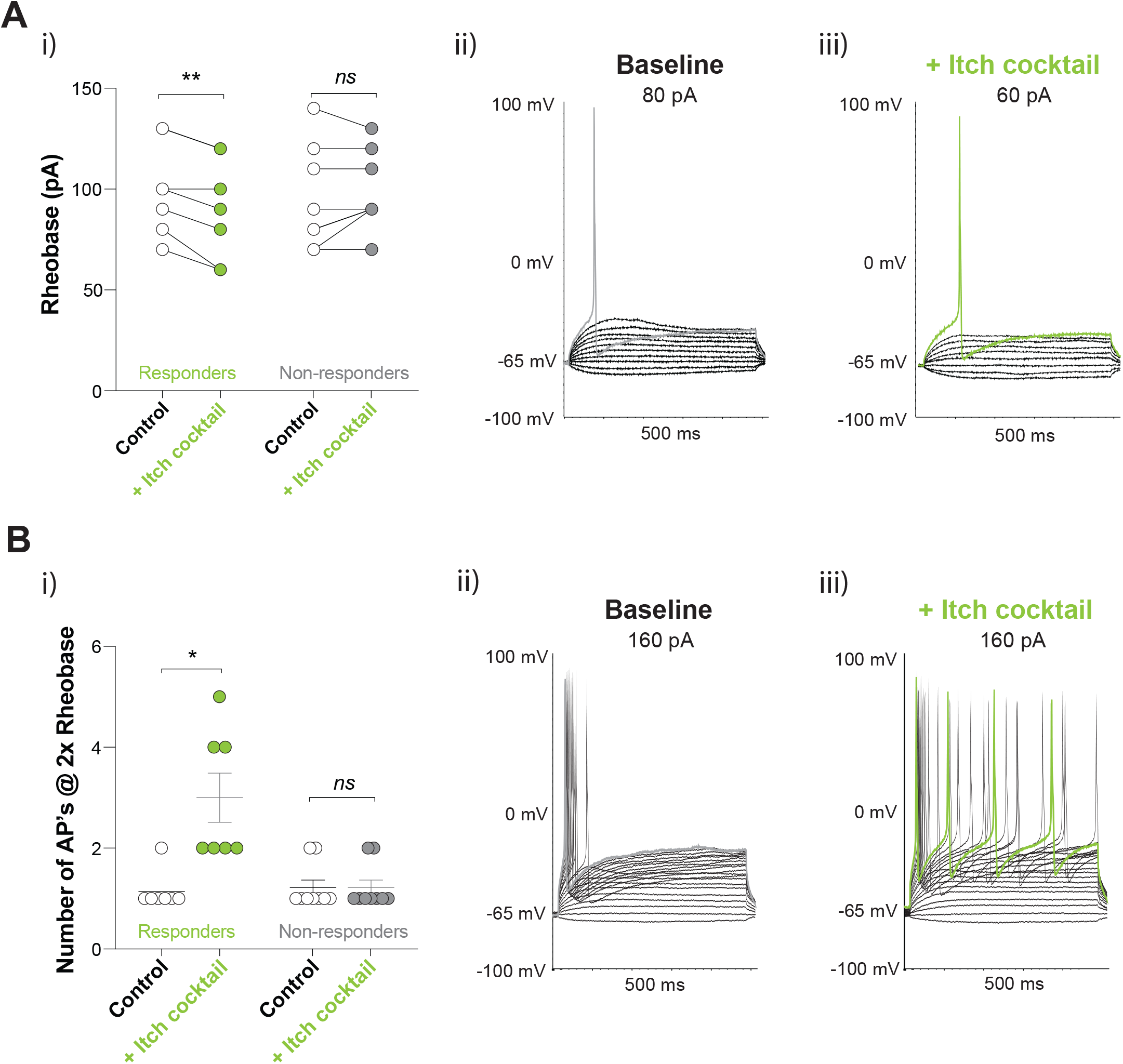
MRGPR agonists induce neuronal hyperexcitability in a cohort of bladder-innervating DRG neurons. **A)** Whole-cell patch clamp recordings of retrogradely traced bladder-innervating LS DRG neurons. **(i)**The Mrgpr agonist itch cocktail significantly decreases the amount of current required to elicit an action potential in a subset (7/16) of bladder-innervating DRG neurons (rheobase; \*\**P*<0.01, paired t test, n=16 neurons from N=5 mice). Another population of bladder-innervating DRG neurons (9/16) were unaffected by the itch cocktail (*ns*, *P*>0.05; paired Student t test, n=9 neurons from N=5 mice). **(ii)** Original recording of membrane voltage from a single bladder-innervating DRG neuron during current ramp protocols. Each line represents a 10pA increase in current. In this example, an 80pA current was required to elicit an action potential under control conditions (grey trace). **(iii)** Perfusion with the ‘itch cocktail’ consisting of chloroquine (1μM), NPFF (5μM) and BAM8-22 (2μM) for 2 minutes onto the same DRG neuron reduced the amount current required to elicit an action potential (green trace), indicating increased neuronal excitability in this ‘responding’ neuron. **B) (i)** Application of the itch cocktail significantly increases the number of action potentials fired at a current twice (2x) the rheobase in a subset (7/16; 44%) of bladder-innervating DRG neurons (\**P*<0.05, Wilcoxon Matched-pairs signed rank test, n=7 neurons from N=5 mice). Other bladder-innervating DRG neurons were unaffected by the itch cocktail (9/16; 56%, ns *P*>0.05, paired t test, n=16 neurons from N=5 mice). **(ii)** Original whole-cell current-clamp recordings from a single bladder-innervating DRG neuron during current clamp protocols. Each line represents a 10pA increase in current. In this example an 80 pA current was required to elicit an action potential under control conditions, with 160 pA representing 2x rheobase (grey trace). **(iii)** Following a 2-minute perfusion with the itch cocktail, the same DRG neuron dramatically increased the number of action potentials at 2x rheobase (green trace), indicating an increase in neuronal excitability. Data in Bi), represent Mean ± SEM. DRG: dorsal root ganglia; LS: lumbosacral; AP’s: Action potentials.

### Peripheral activation of Mrgprs in the bladder results in activation of dorsal horn neurons within the lumbosacral spinal cord

Based on our findings that activation of MrgprA3 and MrgprC11 sensitised the response of a proportion of bladder afferents *ex vivo*, we hypothesised that *in vivo* intra-bladder infusion of Mrgpr agonists should correspondingly enhance bladder signalling into the dorsal horn of the spinal cord. Infusion of saline into the bladder resulted in pERK-immunoreactivity (pERK-IR) in neurons within the LS spinal cord dorsal horn, due to low-level bladder distension **(Figure 8A, 8B)**. In LS spinal segments pERK-IR was seen in the superficial dorsal horn (SDH; **Figure 8Aii, 8B)**, the sacral parasympathetic nucleus (SPN; **Figure 8Aiii, 8B)** and the dorsal grey commissure (DGC; **Figure 8Aiv, 8B)**. Infusion of the ‘itch cocktail’ by co-applying chloroquine, BAM8-22 and NPFF into the bladder resulted in significantly greater total numbers of pERK-IR neurons in the LS spinal cord relative to saline infusion **(Figure 8Ai, 8B)**. Furthermore, significant increases in pERK-IR neurons were observed across the SDH **(Figure 8Aii, 8B)**, SPN **(Figure 8Aiii, 8B)** and the DGC **(Figure 8Aiv, 8B)**, indicating that Mrgpr-evoked bladder afferent hypersensitivity is relayed into the spinal cord and enhances sensory signalling. To confirm that the effects we observed with Mrgpr agonists are mediated via Mrgpr activation we performed the same *in vivo* experiments with *Mrgpr-clusterΔ*^*−/−*^ mice. Comparing Mrgpr agonist and vehicle treated *Mrgpr-clusterΔ*^*−/−*^ mice, we found no differences in the total number of pERK-IR neurons in the LS spinal cord **(Figure 8Ci, 8D)**, the SDH **(Figure 8Cii, 8D)**, the SPN **(Figure 8Ciii, 8D)**, with a small but significant increase in the DGC **(Figure 8Civ, 8D)**.

**Figure 8:**
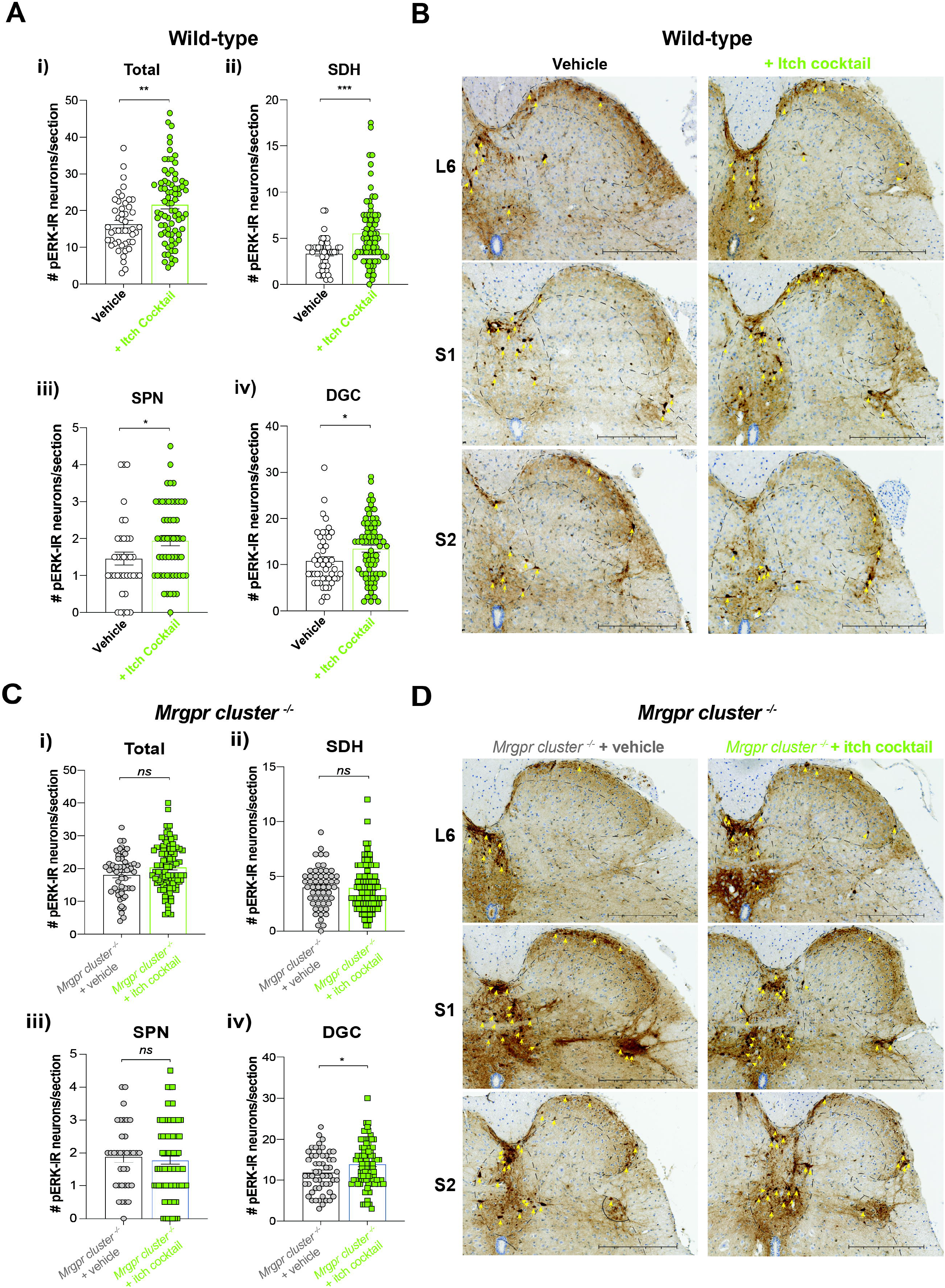
*In vivo* intra-bladder administration of an MRGPR agonist ‘itch cocktail’ results in enhanced activation of dorsal horn neurons within the lumbosacral spinal cord. **A)** Phosphorylated MAPK ERK 1/2 immunoreactivity (pERK-IR) was measured to determine signalling within the lumbosacral (LS; L6–S2) spinal cord in response to intra-bladder administration of saline or an ‘itch cocktail’ of MRGPR agonists (chloroquine: 10μM, BAM8-22: 20μM, NPFF: 5μM). **i)** pERK-IR was significantly increased in the LS dorsal horn of wild-type mice administered the itch cocktail compared with those administered saline (\*\**P*<0.01, Mann Whitney test, n=47-76 sections from spinal regions L6-S2; N=5 mice). ii-iv) Wild-type mice administered the intra-bladder itch cocktail displayed significantly increased pERK-IR in specific regions of the dorsal horn including the **ii)** superficial dorsal horn (SDH), **iii)** sacral parasympathetic nucleus (SPN) and the iv) dorsal gray commissure (DGC) (\**P*<0.05, \*\*\**P*<0.001, Mann Whitney tests). Data are presented as numbers of pERK-IR neurons/section of the LS spinal cord, with a minimum of 6 sections/mouse. **B)** Representative pERK-labelled LS spinal cord images at each spinal level (L6-S2) from mice that had bladders infused with either saline or an MRGPR agonist ‘itch cocktail’. Yellow arrows indicate pERK-IR neurons. Scale bar: 250 μm. **C) i)** In contrast to wild-type mice, pERK-IR was not significantly different across the whole LS dorsal horn of *Mrgpr-clusterΔ*^*−/−*^ mice administered either intra-bladder saline or the MRGPR agonist ‘itch cocktail’ (*ns*, *P*>0.05, Mann Whitney test, n=55-91 sections from N=5 mice). **(ii-iii)** pERK-IR was not significantly altered in the superficial dorsal horn (SDH), **(iii)** nor the sacral parasympathetic nucleus (SPN, *ns*, *P*>0.05, Mann Whitney tests, N=5 mice). iv) Following the ‘itch cocktail’ infusion there was a small but significant increase in pERK-IR in the dorsal grey commissure (DGC) that remained in *Mrgpr-clusterΔ*^*−/−*^ mice (\**P*<0.05; Mann Whitney tests, N=5 mice). Data are presented as numbers of pERK-IR neurons/section of the LS spinal cord, with a minimum of 6 sections/mouse. **D)** Representative pERK-labelled LS spinal cord images at each spinal level (L6-S1) from *Mrgpr-clusterΔ*^*−/−*^ mice that had bladders infused with either saline or the MRGPR agonist ‘itch cocktail’. Yellow arrows indicate pERK-IR neurons. Scale bars: 250 μm. Data represent Mean ± SEM

### Baseline bladder afferent responses and signalling within the spinal cord to bladder distension are unaltered in *Mrgpr-clusterΔ*^*−/−*^ mice

In order to determine if Mrgpr’s contribute to bladder afferent responses to distension at baseline, we compared responses in wildtype and *Mrgpr-clusterΔ*^*−/−*^ littermate mice. *Ex vivo* bladder afferent firing to bladder distension with saline showed that responses were indistinguishable between wildtype and *Mrgpr-clusterΔ*^*−/−*^ mice **(Figure 9A)**. Similarly, following saline bladder distension *in vivo* the numbers of pERK-IR neurons within the LS spinal cord were not significantly different between wild-type and *Mrgpr-clusterΔ*^*−/−*^ mice **(Figure 9B, 9C, 9D, 9E)**. Overall, these findings suggest Mrgprs are not responsible for bladder afferent mechanosensitivity *per se*, rather have a prominent role in sensitizing bladder afferent responsiveness to distension following activation by pruritogens.

**Figure 9:**
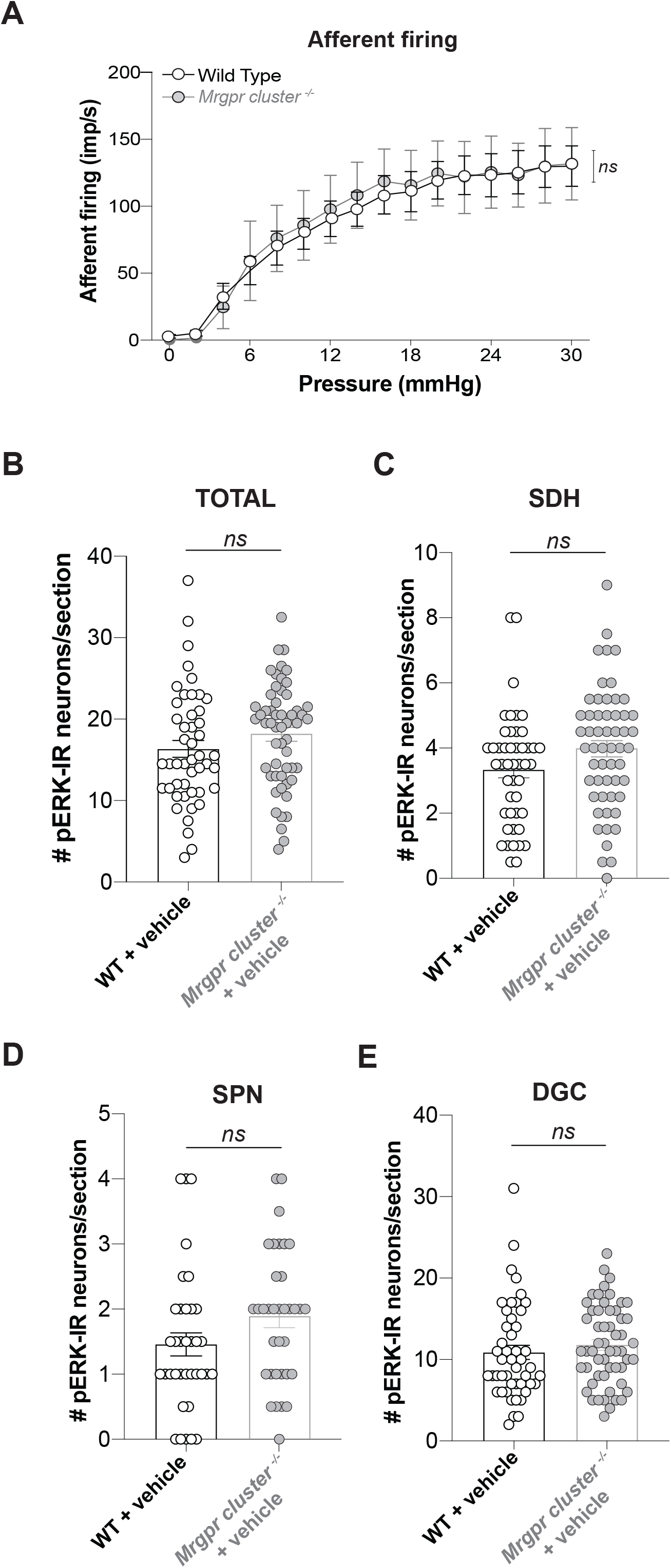
*Mrgpr-clusterΔ*^*−/−*^ mice do not display altered baseline responses to bladder distension. **A)** Group data from *ex vivo* recordings from wild-type mice and age matched *Mrgpr-clusterΔ*^*−/−*^ mice. There was no significant difference in the mechanosensitivity of bladder afferents in response to graded bladder distension (0-30mmHg) between wild-type and *Mrgpr-clusterΔ*^*−/−*^ mice (*ns*, *P*>0.05, 2-way ANOVA, N=4-15 mice). **B-E)** Compared with wild-type mice *Mrgpr-clusterΔ*^*−/−*^ mice do not display altered neuronal activation within the dorsal horn of LS spinal cord following intra-bladder vehicle administration. Phosphorylated MAPK ERK 1/2 immunoreactivity (pERK-IR) was measured to determine signalling in the lumbosacral (LS; L6–S2) spinal cord in response to saline instillation in wildtype and age matched *Mrgpr-clusterΔ*^*−/−*^ mice. There was no difference in the (**B)** total number of pERK-IR neurons identified within the LS dorsal horn in response to saline infusion into the bladder between wildtype and *Mrgpr-clusterΔ*^*−/−*^ mice. In terms of specific regions of the LS dorsal horn, there were significant differences between wild-type and *Mrgpr-clusterΔ*^*−/−*^ mice in the number of pERK-IR neurons following intra-bladder administration within the **(C)** SDH, **(D)** SPN and **(E)** DGC regions of the dorsal horn (*ns*, *P*>0.05, Mann Whitney tests, N=5 mice/group). Lumbosacral: LS. Data represent Mean ± SEM.

## DISCUSSION

The ability of the somatosensory nervous system to easily differentiate stimuli such as pruritogens or algogens is essential as it evokes a tailored response, such as scratching or withdrawal of a limb. As a consequence, there is significant functional and anatomical diversity of somatosensory afferent nerves. In contrast, sensory discrimination in the bladder is more limited in scope, ranging from fullness to urge and finally pain (de Groat et al., 2015). Therefore, the differentiation of distinct sensory stimuli (inflammation, infection, irritation) is less important if the overarching physiological response remains the same – increased voiding to rapidly remove irritants from the bladder. As such, mechanosensitive afferents innervating the bladder are considerably less diverse than cutaneous afferents. The majority of bladder neurons are characterised as peptidergic C-fibers that express *Trpv1* (de Groat et al., 2015; Grundy et al., 2019b), are embedded within the detrusor smooth muscle (de Groat et al., 2015; Grundy et al., 2019a; Grundy et al., 2019b), and are broadly tuned to allow the detection of a variety of mechanical and chemical stimuli (Rong et al., 2002; Zagorodnyuk et al., 2006; Zagorodnyuk et al., 2007). Therefore, the excitability of these afferents is key in providing appropriate sensory information in response to stimuli, including bladder filling and emptying. Understanding the mechanisms responsible for regulating bladder afferent mechanosensitivity is therefore crucial to the development of novel treatments for the chronic sensory symptoms associated with bladder hypersensitivity and dysfunction, including urinary urgency, frequency and pelvic pain. In this study we discovered a novel role in bladder sensory afferent pathways for irritant sensing mechanisms mediated via MrgprC11 and MrgprA3.

Consistent with previous reports that Mrgpr agonists are able to excite isolated peripheral sensory neurons (Liu et al., 2009; Castro et al., 2019; Tseng et al., 2019) we showed, using patch clamp recordings, that chloroquine, BAM8-22 and NPFF activated and induced neuronal hyperexcitability in sub-populations (44%) of bladder-innervating DRG neurons. In calcium imaging experiments, we observed some functional overlap between responses to chloroquine and BAM8-22, which has been previously described in afferents innervating the skin (Liu et al., 2009) and colon (Castro et al., 2019). Employing single-cell RT-PCR of bladder DRG neurons, we revealed expression and co-expression of Mrgprs in bladder-innervating neurons. The majority of neurons expressing *MrgprC11* also contained *MrgprA3*, consistent with previous reports of colocalization in other sensory pathways (Dong et al., 2001; Zylka et al., 2003; Han et al., 2013; Castro et al., 2019).

High colocalization of *MrgprA3* and *MrgprC11* was also observed with *Trpv*1 in bladder DRG neurons, either on an mRNA level with single cell RT-PCR, or functionally with bladder neurons exhibiting calcium transients in response to chloroquine, BAM8-22, NPFF and the TRPV1 agonist capsaicin. This is expected as the majority of bladder-innervating DRG neurons express *Trpv1* (La et al., 2011; Dang et al., 2013; Forrest et al., 2013), whilst TRPV1 is a key integrator of histamine and P2X_3_ dependent bladder afferent activation (Grundy et al., 2018a; Grundy et al., 2020). These findings are consistent with the literature suggesting *MrgprA3*-positive neurons are in a sub-population of *Trpv1*-positive neurons (Han et al., 2013; Usoskin et al., 2015; Meixiong and Dong, 2017). However, these mechanisms in the bladder, where only 15% of bladder neurons express *Trpa1*, appear to contrast with those in the colon and skin. For example, Mrgpr agonist-induced mechanical hypersensitivity to colorectal distension is not evoked in *Trpa1*^*−/−*^ mice (Castro et al., 2019). In the skin TRPA1 has been identified as the downstream target of both MrgprA3 and MrgprC11, whereby *Trpa1*^*−/−*^ mice display little to no scratching in response to chloroquine or BAM8-22, with functional coupling of MrgprA3 or MrgprC11 to TRPA1 occurring via Gβγ and PLC signalling respectively (Wilson et al., 2011).

Differences are also apparent between bladder and cutaneous neurons when interrogating the role of MrgprD. Within cutaneous populations two distinct pruritogenic sub-populations of DRG neurons have been identified; those expressing *MrgprA3, MrgprC11* and histamine receptor 1 (*H*_*1*_*R*), and those expressing *MrgprD* (Liu et al., 2009; Liu et al., 2012; Usoskin et al., 2015). MrgprD, initially postulated to play an important nociceptive role, is now known to mediate itch induced by the amino acid β-alanine (Shinohara et al., 2004; Liu et al., 2012). Despite high MrgprD expression in whole LS DRG, only 6% of bladder-innervating DRG neurons expressed MrgprD, which showed equal co-expression (40%) with MrgprC11 and MrgprA3. This suggests a far more prominent role for MrgprC11 and MrgprA3 than MrgprD in bladder sensory function. In keeping with this reasoning, we show that chloroquine, BAM8-22 and NPFF enhanced the mechanosensitivity of sub-populations (20-41%) of wild-type bladder afferents to bladder distension, which did not occur in experiments with *Mrgpr-clusterΔ*^*−/−*^ mice. The proportion of wild-type afferents responding to these Mrgpr agonists correlated well with the proportion of bladder neurons expressing MrgprA3 (14%) and MrgprC11 (38%). Mrgpr mRNA expression in bladder tissue was extremely low or below the detection limit, explaining why intra-bladder instillation of Mrgpr agonists did not affect bladder compliance. These findings are in keeping with a number of studies showing that these Mrgprs are expressed almost exclusively in sensory structures, including cell bodies of sensory neurons located in the DRG, trigeminal and jugular ganglia (Dong et al., 2001; Han et al., 2013; Meixiong and Dong, 2017; Han et al., 2018; Castro et al., 2019).

We found that Mrgpr agonists evoked bladder afferent mechanical hypersensitivity across a broad range of distension pressures. Unlike other irritant and inflammatory stimuli, such as histamine or inflammatory cytokines, Mrgpr agonist incubation did not recruit silent afferents to become mechanosensitive (Brierley et al., 2020; Grundy et al., 2020). This is an important distinction as silent afferents represent a discrete functional subclass of bladder afferents, that after becoming sensitized acquire an ability to respond to ‘high-threshold’ or ‘nociceptive’ mechanosensory stimuli (Prato et al., 2017; Zhang et al., 2019; Brierley et al., 2020). These findings are consistent with MrgprA3- and MrgprC11-positive neurons that have previously been reported as being responsive to mechanical stimuli, including probing of cutaneous receptive fields (Han et al., 2013). Furthermore, a recent study investigating the role of Mrgprs in colonic afferent mechanosensitivity found that MrgprA3- and MrgprC11-positive fibres are also sensitive to mechanical stimulation (Castro et al., 2019). Mrgpr-positive cutaneous nerve endings have been reported to exclusively innervate the epidermis (Han et al., 2013), however this does not appear to be the case in the bladder. Whilst some mechanosensitive afferents innervate the bladder mucosa, an equivalent to the epidermis, the majority of mechanosensitive bladder afferents are found innervating the bladder detrusor (Zagorodnyuk et al., 2007; Spencer et al., 2018), and we observed significant hypersensitivity of these distension sensitive afferents to Mrgpr agonists.

The homogeneity of bladder sensory pathways is further reflected in their activation of spinal cord circuits in response to bladder distension, which unlike the distinct organisation of primary afferent circuits that occurs in the spinal cord for processing specific somatosensory stimuli (Todd, 2010), appears graded only to the level of peripheral stimuli (Grundy et al., 2018a). As such, an increase in the intensity of the peripheral afferent signal corresponds to increased activation of second-order neurons that project within multiple visceral sensory pathways to regulate networks responsible for both micturition and sensation, rather than any obvious sensation-specific activation. Accordingly, we found Mrgpr activation of peripheral afferents in the bladder *in vivo* resulted in significant increases in the number of pERK-immunoreactive neurons in all regions of the lumbosacral dorsal horn that receive input from mechanosensitive afferents (Grundy et al., 2019b). This included increased numbers of neurons within the superficial dorsal horn, dorsal grey commissure and sacral parasympathetic nucleus (SPN). As such, we would expect that the physiological consequence of an Mrgpr-induced increase in afferent firing would be similar to other stimuli that evoke an increase in primary afferent excitability from the bladder - urinary urgency and urinary frequency.

Interestingly, we observed no difference in bladder afferent mechanosensitivity nor activation of central pathways in the spinal cord to distension with saline between *Mrgpr-clusterΔ*^*−/−*^ mice and wild-type mice. Similar observations have been made before in other sensory pathways, where *Mrgpr-clusterΔ*^*−/−*^ mice do not show deficits in cutaneous mechanosensation (Liu et al., 2009). Furthermore, in the colon *Mrgpr-clusterΔ*^*−/−*^ mice had similar visceromotor responses to wild-type mice (Castro et al., 2019). Overall, these data suggest that endogenous activators of MrgprC11 and MrgprA3 are not tonically released from the bladder to set the baseline tone of bladder afferent mechanosensitivity. Whether or not these mechanisms are up-regulated in conditions of bladder pathophysiology is the subject of future research.

In conclusion, we have shown that MrgprC11 and MrgprA3 are present and functional on mechanosensitive bladder afferents and their activation induces mechanical hypersensitivity and increased activation of sensory circuits within the spinal cord. Enhanced sensation via an increase in mechanosensitivity may be a fundamental protective mechanism and relate more specifically to bladder disorders associated with urgency and frequency in the absence of pain, such as OAB.

## Acknowledgements

Work was supported by a National Health and Medical Research Council of Australia (NHMRC) Project Grant (APP1140297 to S.M.B), an NHMRC R.D Wright Biomedical Research Fellowship (APP1126378 to S.M.B) and an Australian Research Council (ARC) Discovery Project (DP180101395 to A.M.H and S.M.B).

## REFERENCES

Abrams P, Cardozo L, Fall M, Griffiths D, Rosier P, Ulmsten U, van Kerrebroeck P, Victor A, Wein A, Standardisation Sub-committee of the International Continence S (2002) The standardisation of terminology of lower urinary tract function: report from the Standardisation Sub-committee of the International Continence Society. Neurourol Urodyn 21:167–178.

Andersson KE (2019) TRP Channels as Lower Urinary Tract Sensory Targets. Med Sci (Basel) 7:67.

Bellono NW, Bayrer JR, Leitch DB, Castro J, Zhang C, O’Donnell TA, Brierley SM, Ingraham HA, Julius D (2017) Enterochromaffin Cells Are Gut Chemosensors that Couple to Sensory Neural Pathways. Cell 170:185–198 e116.

Brierley SM, Jones RC, 3rd, Gebhart GF, Blackshaw LA (2004) Splanchnic and pelvic mechanosensory afferents signal different qualities of colonic stimuli in mice. Gastroenterology 127:166–178.

Brierley SM, Goh KGK, Sullivan MJ, Moore KH, Ulett GC, Grundy L (2020) Innate immune response to bacterial urinary tract infection sensitises high-threshold bladder afferents and recruits silent nociceptors. Pain 161:202–210.

Castro J, Harrington AM, Garcia-Caraballo S, Maddern J, Grundy L, Zhang J, Page G, Miller PE, Craik DJ, Adams DJ, Brierley SM (2017) alpha-Conotoxin Vc1.1 inhibits human dorsal root ganglion neuroexcitability and mouse colonic nociception via GABAB receptors. Gut 66:1083–1094.

Castro J, Harrington AM, Lieu T, Garcia-Caraballo S, Maddern J, Schober G, O’Donnell T, Grundy L, Lumsden AL, Miller P, Ghetti A, Steinhoff MS, Poole DP, Dong X, Chang L, Bunnett NW, Brierley SM (2019) Activation of pruritogenic TGR5, MrgprA3, and MrgprC11 on colon-innervating afferents induces visceral hypersensitivity. JCI Insight 4.

Christianson JA, Liang R, Ustinova EE, Davis BM, Fraser MO, Pezzone MA (2007) Convergence of bladder and colon sensory innervation occurs at the primary afferent level. Pain 128:235–243.

Daly D, Rong W, Chess-Williams R, Chapple C, Grundy D (2007) Bladder afferent sensitivity in wild-type and TRPV1 knockout mice. The Journal of physiology 583:663–674.

Dang K, Bielefeldt K, Gebhart GF (2013) Cyclophosphamide-induced cystitis reduces ASIC channel but enhances TRPV1 receptor function in rat bladder sensory neurons. J Neurophysiol 110:408–417.

de Groat WC, Yoshimura N (2009) Afferent nerve regulation of bladder function in health and disease. Handb Exp Pharmacol:91–138.

de Groat WC, Griffiths D, Yoshimura N (2015) Neural control of the lower urinary tract. Compr Physiol 5:327–396.

DeBerry JJ, Schwartz ES, Davis BM (2014) TRPA1 mediates bladder hyperalgesia in a mouse model of cystitis. Pain 155:1280–1287.

Dong X, Han S, Zylka MJ, Simon MI, Anderson DJ (2001) A diverse family of GPCRs expressed in specific subsets of nociceptive sensory neurons. Cell 106:619–632.

Everaerts W, Vriens J, Owsianik G, Appendino G, Voets T, De Ridder D, Nilius B (2010a) Functional characterization of transient receptor potential channels in mouse urothelial cells. American journal of physiology Renal physiology 298:F692–701.

Everaerts W, Zhen X, Ghosh D, Vriens J, Gevaert T, Gilbert JP, Hayward NJ, McNamara CR, Xue F, Moran MM, Strassmaier T, Uykal E, Owsianik G, Vennekens R, De Ridder D, Nilius B, Fanger CM, Voets T (2010b) Inhibition of the cation channel TRPV4 improves bladder function in mice and rats with cyclophosphamide-induced cystitis. Proc Natl Acad Sci U S A 107:19084–19089.

Forrest SL, Osborne PB, Keast JR (2013) Characterization of bladder sensory neurons in the context of myelination, receptors for pain modulators, and acute responses to bladder inflammation. Frontiers in neuroscience 7:206.

Gao YJ, Ji RR (2009) c-Fos and pERK, which is a better marker for neuronal activation and central sensitization after noxious stimulation and tissue injury? Open Pain J 2:11–17.

Grundy L, Caldwell A, Brierley SM (2018a) Mechanisms Underlying Overactive Bladder and Interstitial Cystitis/Painful Bladder Syndrome. Frontiers in neuroscience 12:931.

Grundy L, Erickson A, Brierley SM (2019a) Visceral Pain. Annu Rev Physiol 81:261–284.

Grundy L, Daly DM, Chapple C, Grundy D, Chess-Williams R (2018b) TRPV1 enhances the afferent response to P2X receptor activation in the mouse urinary bladder. Sci Rep 8:197.

Grundy L, Erickson A, Caldwell A, Garcia-Caraballo S, Rychkov G, Harrington A, Brierley SM (2018c) Tetrodotoxin-sensitive voltage-gated sodium channels regulate bladder afferent responses to distension. Pain 159:2573–2584.

Grundy L, Caldwell A, Garcia Caraballo S, Erickson A, Schober G, Castro J, Harrington AM, Brierley SM (2020) Histamine induces peripheral and central hypersensitivity to bladder distension via the histamine H1 receptor and TRPV1. American journal of physiology Renal physiology 318:F298–F314.

Grundy L, Chess-Williams R, Brierley SM, Mills K, Moore KH, Mansfield K, Rose’Meyer R, Sellers D, Grundy D (2018d) NKA enhances bladder-afferent mechanosensitivity via urothelial and detrusor activation. American journal of physiology Renal physiology 315:F1174–F1185.

Grundy L, Harrington AM, Caldwell A, Castro J, Staikopoulos V, Zagorodnyuk VP, Brookes SJH, Spencer NJ, Brierley SM (2019b) Translating peripheral bladder afferent mechanosensitivity to neuronal activation within the lumbosacral spinal cord of mice. Pain 160:793–804.

Grundy L, Harrington AM, Castro J, Garcia-Caraballo S, Deiteren A, Maddern J, Rychkov GY, Ge P, Peters S, Feil R, Miller P, Ghetti A, Hannig G, Kurtz CB, Silos-Santiago I, Brierley SM (2018e) Chronic linaclotide treatment reduces colitis-induced neuroplasticity and reverses persistent bladder dysfunction. JCI Insight 3.

Han L, Limjunyawong N, Ru F, Li Z, Hall OJ, Steele H, Zhu Y, Wilson J, Mitzner W, Kollarik M, Undem BJ, Canning BJ, Dong X (2018) Mrgprs on vagal sensory neurons contribute to bronchoconstriction and airway hyper-responsiveness. Nat Neurosci 21:324–328.

Han L, Ma C, Liu Q, Weng HJ, Cui Y, Tang Z, Kim Y, Nie H, Qu L, Patel KN, Li Z, McNeil B, He S, Guan Y, Xiao B, Lamotte RH, Dong X (2013) A subpopulation of nociceptors specifically linked to itch. Nat Neurosci 16:174–182.

Harrington AM, Caraballo SG, Maddern JE, Grundy L, Castro J, Brierley SM (2019) Colonic afferent input and dorsal horn neuron activation differs between the thoracolumbar and lumbosacral spinal cord. Am J Physiol Gastrointest Liver Physiol 317:G285–G303.

Homma Y, Akiyama Y, Tomoe H, Furuta A, Ueda T, Maeda D, Lin AT, Kuo HC, Lee MH, Oh SJ, Kim JC, Lee KS (2020) Clinical guidelines for interstitial cystitis/bladder pain syndrome. Int J Urol 27:578–589.

Inserra MC, Israel MR, Caldwell A, Castro J, Deuis JR, Harrington AM, Keramidas A, Garcia-Caraballo S, Maddern J, Erickson A, Grundy L, Rychkov GY, Zimmermann K, Lewis RJ, Brierley SM, Vetter I (2017) Multiple sodium channel isoforms mediate the pathological effects of Pacific ciguatoxin-1. Sci Rep 7:42810.

Jiang GY, Dai MH, Huang K, Chai GD, Chen JY, Chen L, Lang B, Wang QX, St Clair D, McCaig C, Ding YQ, Zhang L (2015) Neurochemical characterization of pERK-expressing spinal neurons in histamine-induced itch. Sci Rep 5:12787.

Konthapakdee N, Grundy L, O’Donnell T, Garcia-Caraballo S, Brierley SM, Grundy D, Daly DM (2019) Serotonin exerts a direct modulatory role on bladder afferent firing in mice. The Journal of physiology 597:5247–5264.

Kreis ME, Haupt W, Kirkup AJ, Grundy D (1998) Histamine sensitivity of mesenteric afferent nerves in the rat jejunum. Am J Physiol 275:G675–680.

La JH, Schwartz ES, Gebhart GF (2011) Differences in the expression of transient receptor potential channel V1, transient receptor potential channel A1 and mechanosensitive two pore-domain K+ channels between the lumbar splanchnic and pelvic nerve innervations of mouse urinary bladder and colon. Neuroscience 186:179–187.

LaMotte RH, Dong X, Ringkamp M (2014) Sensory neurons and circuits mediating itch. Nat Rev Neurosci 15:19–31.

Lee JS, Han JS, Lee K, Bang J, Lee H (2016) The peripheral and central mechanisms underlying itch. BMB Rep 49:474–487.

Lee MG, Dong X, Liu Q, Patel KN, Choi OH, Vonakis B, Undem BJ (2008) Agonists of the MAS-related gene (Mrgs) orphan receptors as novel mediators of mast cell-sensory nerve interactions. J Immunol 180:2251–2255.

Lein ES et al. (2007) Genome-wide atlas of gene expression in the adult mouse brain. Nature 445:168–176.

Lembo PM, Grazzini E, Groblewski T, O’Donnell D, Roy MO, Zhang J, Hoffert C, Cao J, Schmidt R, Pelletier M, Labarre M, Gosselin M, Fortin Y, Banville D, Shen SH, Strom P, Payza K, Dray A, Walker P, Ahmad S (2002) Proenkephalin A gene products activate a new family of sensory neuron--specific GPCRs. Nat Neurosci 5:201–209.

Liu Q, Sikand P, Ma C, Tang Z, Han L, Li Z, Sun S, LaMotte RH, Dong X (2012) Mechanisms of itch evoked by beta-alanine. J Neurosci 32:14532–14537.

Liu Q, Tang Z, Surdenikova L, Kim S, Patel KN, Kim A, Ru F, Guan Y, Weng HJ, Geng Y, Undem BJ, Kollarik M, Chen ZF, Anderson DJ, Dong X (2009) Sensory neuron-specific GPCR Mrgprs are itch receptors mediating chloroquine-induced pruritus. Cell 139:1353–1365.

Liu T, Ji RR (2013) New insights into the mechanisms of itch: are pain and itch controlled by distinct mechanisms? Pflugers Arch 465:1671–1685.

Meixiong J, Dong X (2017) Mas-Related G Protein-Coupled Receptors and the Biology of Itch Sensation. Annu Rev Genet 51:103–121.

Osteen JD, Herzig V, Gilchrist J, Emrick JJ, Zhang C, Wang X, Castro J, Garcia-Caraballo S, Grundy L, Rychkov GY, Weyer AD, Dekan Z, Undheim EA, Alewood P, Stucky CL, Brierley SM, Basbaum AI, Bosmans F, King GF, Julius D (2016) Selective spider toxins reveal a role for the Nav1.1 channel in mechanical pain. Nature 534:494–499.

Prato V, Taberner FJ, Hockley JRF, Callejo G, Arcourt A, Tazir B, Hammer L, Schad P, Heppenstall PA, Smith ES, Lechner SG (2017) Functional and Molecular Characterization of Mechanoinsensitive “Silent” Nociceptors. Cell Rep 21:3102–3115.

Rong W, Spyer KM, Burnstock G (2002) Activation and sensitisation of low and high threshold afferent fibres mediated by P2X receptors in the mouse urinary bladder. The Journal of physiology 541:591–600.

Schmelz M (2015) Neurophysiology and itch pathways. Handb Exp Pharmacol 226:39–55.

Shim WS, Oh U (2008) Histamine-induced itch and its relationship with pain. Mol Pain 4:29.

Shinohara T, Harada M, Ogi K, Maruyama M, Fujii R, Tanaka H, Fukusumi S, Komatsu H, Hosoya M, Noguchi Y, Watanabe T, Moriya T, Itoh Y, Hinuma S (2004) Identification of a G protein-coupled receptor specifically responsive to beta-alanine. J Biol Chem 279:23559–23564.

Sikand P, Dong X, LaMotte RH (2011) BAM8-22 peptide produces itch and nociceptive sensations in humans independent of histamine release. J Neurosci 31:7563–7567.

Spencer NJ, Greenheigh S, Kyloh M, Hibberd TJ, Sharma H, Grundy L, Brierley SM, Harrington AM, Beckett EA, Brookes SJ, Zagorodnyuk VP (2018) Identifying unique subtypes of spinal afferent nerve endings within the urinary bladder of mice. J Comp Neurol 526:707–720.

Steinhoff M, Schmelz M, Szabo IL, Oaklander AL (2018) Clinical presentation, management, and pathophysiology of neuropathic itch. The Lancet Neurology 17:709–720.

Todd AJ (2010) Neuronal circuitry for pain processing in the dorsal horn. Nat Rev Neurosci 11:823–836.

Tominaga M, Takamori K (2014) Sensitization of Itch Signaling: Itch Sensitization-Nerve Growth Factor, Semaphorins. In: Itch: Mechanisms and Treatment (Carstens E, Akiyama T, eds). Boca Raton (FL): CRC Press/Taylor & Francis (c) 2014 by Taylor & Francis Group, LLC.

Tseng P-Y, Zheng Q, Li Z, Dong X (2019) MrgprX1 mediates neuronal excitability and itch through tetrodotoxin-resistant sodium channels. Itch 4:e28.

Usoskin D, Furlan A, Islam S, Abdo H, Lonnerberg P, Lou D, Hjerling-Leffler J, Haeggstrom J, Kharchenko O, Kharchenko PV, Linnarsson S, Ernfors P (2015) Unbiased classification of sensory neuron types by large-scale single-cell RNA sequencing. Nat Neurosci 18:145–153.

Van Remoortel S, Ceuleers H, Arora R, Van Nassauw L, De Man JG, Buckinx R, De Winter BY, Timmermans JP (2019) Mas-related G protein-coupled receptor C11 (Mrgprc11) induces visceral hypersensitivity in the mouse colon: A novel target in gut nociception? Neurogastroenterol Motil 31:e13623.

Wilson SR, Gerhold KA, Bifolck-Fisher A, Liu Q, Patel KN, Dong X, Bautista DM (2011) TRPA1 is required for histamine-independent, Mas-related G protein-coupled receptor-mediated itch. Nat Neurosci 14:595–602.

Wouters MM et al. (2016) Histamine Receptor H1-Mediated Sensitization of TRPV1 Mediates Visceral Hypersensitivity and Symptoms in Patients With Irritable Bowel Syndrome. Gastroenterology 150:875–887 e879.

Xu L, Gebhart GF (2008) Characterization of mouse lumbar splanchnic and pelvic nerve urinary bladder mechanosensory afferents. J Neurophysiol 99:244–253.

Yoshimura N, Ogawa T, Miyazato M, Kitta T, Furuta A, Chancellor MB, Tyagi P (2014) Neural mechanisms underlying lower urinary tract dysfunction. Korean J Urol 55:81–90.

Zagorodnyuk VP, Costa M, Brookes SJ (2006) Major classes of sensory neurons to the urinary bladder. Auton Neurosci 126-127:390–397.

Zagorodnyuk VP, Gibbins IL, Costa M, Brookes SJ, Gregory SJ (2007) Properties of the major classes of mechanoreceptors in the guinea pig bladder. The Journal of physiology 585:147–163.

Zhang Y, Li S, Yecies T, Morgan T, Cai H, Pace N, Shen B, Wang J, Roppolo JR, de Groat WC, Tai C (2019) Sympathetic afferents in the hypogastric nerve facilitate nociceptive bladder activity in cats. American journal of physiology Renal physiology 316:F703–F711.

Zhu Y, Hanson CE, Liu Q, Han L (2017) Mrgprs activation is required for chronic itch conditions in mice. Itch (Phila) 2.

Zylka MJ, Dong X, Southwell AL, Anderson DJ (2003) Atypical expansion in mice of the sensory neuron-specific Mrg G protein-coupled receptor family. Proc Natl Acad Sci U S A 100:10043–10048.

